# End-to-end deep learning versus machine learning for biomarker discovery in cancer genomes

**DOI:** 10.1101/2025.01.06.631471

**Authors:** Michaela Unger, Chiara M. L. Loeffler, Julien Vibert, Laura Žigutytė, Srividhya Sainath, Tim Lenz, Andreas Mock, Stefan Fröhling, Trevor A. Graham, Zunamys I. Carrero, Jakob Nikolas Kather

## Abstract

**Background:** Accurate determination of genomic biomarkers from tumor sequencing is fundamental to precision oncology, informing disease classification and treatment decisions. In practice, biomarker inference relies on computational pipelines that often compress high-dimensional mutation data into predefined summaries such as mutational signatures or composite genomic features. While robust and widely adopted, these representations may not fully capture the complexity of cancer genomes. Deep learning (DL) offers an end-to-end alternative by learning features directly from raw genomic data. However, clinical translation remains challenging due to limited empirical validation of new DL models and a lack of systematic comparisons with established machine learning (ML) baselines, particularly when transitioning from information-rich genome or exome data to real-world targeted sequencing profiles. Here, we compare state-of-the-art DL architectures with classical ML models across variant-level, copy-number (CNV), and multimodal inputs, using microsatellite instability (MSI) and homologous recombination deficiency (HRD) prediction as oncologically relevant tasks. We aim to derive practical guidance on modelling strategies across different data modalities and clinical sequencing contexts.

**Methods:** For MSI and HRD prediction, we trained multiple DL models, including supervised and self-supervised encoders, alongside feature-based ML approaches using tumor mutation data, copy-number alterations, and their multimodal combinations. Analyses were conducted on 5,647 patients in The Cancer Genome Atlas (TCGA), the Clinical Proteomic Tumor Analysis Consortium (CPTAC), and two targeted sequencing panel cohorts. Model performance was evaluated on both whole-exome and panel-based datasets, and explainability analysis were performed for both DL and ML models.

**Results:** For MSI, DL demonstrated stronger generalization than ML on external validation data (F1 0.97 vs 0.76) and maintained comparatively high performance under pseudo-panels conditions, whereas ML performance dropped. In a real-world targeted panel cohort, DL again showed more robust generalization than ML, with performance partly affected by cross-assay variability. For HRD, incorporation of CNV data was the primary determinant of predictive performance. Once CNVs were included, DL and ML achieved similar accuracy on external datasets (F1 0.61 vs 0.58). In panel-based settings, DL retained an advantage over ML (F1 0.78 vs 0.62). Model interpretation analyses indicated that both DL and ML relied on mutation and chromosomal patterns consistent with established MSI and HRD biology.

**Conclusion:** Overall, predictive performance depended strongly on data availability and clinical sequencing context. When information-rich inputs were available, both DL and classical ML achieved robust biomarker prediction, with DL generally matching or exceeding ML performance. The most pronounced advantages of DL emerged in cross-assay evaluations and data-sparse settings, where generalization was more reliable. Notably, the best-performing DL models were lightweight and interpretable, supporting practical deployment. In clinical genomics workflows, such models may complement established pipelines by leveraging patient sequencing data to provide additional evidence for treatment-relevant biomarker assessment.

## Background

Bulk and targeted sequencing of patient tumors are being progressively integrated into routine clinical workflows. Genomic information has become a central component of personalized cancer care, helping to refine diagnosis, identify hereditary predisposition, guide systemic treatment choices, and explain therapeutic resistance [1–5]. In current practice, next-generation sequencing (NGS) data are processed through multi-step bioinformatic pipelines designed around predefined genomic features. Information from tumor sequencing is often summarized into hand-crafted features such as curated gene lists, mutational and structural signatures, or other aggregated metrics [6–11]. While these approaches are well established, they inevitably rely on extensive a priori assumptions and may not fully exploit potentially relevant genomic signals contained in raw data. This limitation becomes more pronounced as profiling moves from genome- and exome-wide assays to targeted panels, where fewer observed events must support the same clinical decisions.

Artificial intelligence (AI) has been proposed as a way to reduce this dependency on manually engineered features. In recent years, deep learning (DL) models have been increasingly applied to patient-level genomic prediction tasks as sequencing costs decline and computational resources increase [12–16]. In principle, DL can learn task-relevant features directly from raw genomic inputs, potentially capturing mutational patterns that are difficult to specify a priori and that may improve upon traditional representations. At the same time, many DL studies in genomics remain difficult to translate into clinical practice because they are not systematically benchmarked against strong classical machine learning (ML) baselines, and because evaluation on independent cohorts is often limited. This has raised concerns about fair benchmarking and the risk of overengineering [17–20]. For clinical adoption, it is therefore crucial to understand when end-to-end DL provides a tangible advantage, and when simpler feature-based pipelines remain sufficient, particularly in the targeted-panel setting that dominates routine testing.

To address these questions in a clinically grounded manner, we use two established compound biomarkers as representative test cases: microsatellite instability (MSI) and homologous recombination deficiency (HRD). Both reflect characteristic forms of genomic instability and have direct therapeutic implications. MSI arises from defects in the mismatch repair (MMR) pathway, leading to an accumulation of single-nucleotide variants (SNVs) and small insertions and deletions (indels), particularly in repetitive regions. Tumours with high MSI are strong candidates for immune checkpoint inhibition [21,22]. HRD, in contrast, reflects impaired repair of DNA double-strand breaks and leaves a characteristic pattern of SNVs, indels, structural variants (SVs), and copy-number variations (CNVs) in the cancer genome. HRD status is associated with sensitivity to PARP inhibitors in several tumor types [23–26]. Thus, both MSI and HRD encode biologically meaningful information directly within the cancer genome in the form of mutational patterns [9]. Therefore, these biomarkers are clinically relevant and provide complementary test cases across sequencing assays and data constraints, making them well suited for benchmarking modeling strategies [27–29].

For MSI prediction, several DL approaches have relied on specialized or non-routine sequencing assays rather than on data generated typically in clinical practice, where MSI testing often begins with immunohistochemistry and NGS is predominantly panel-based [30–32]. For HRD, a range of established predictors exists, including HRDetect [33], CHORD [34], and HRProfiler [35], but these tools depend on curated mutation signatures, selected variants, or complex SV features and tend to perform best on whole-genome sequencing (WGS). Their performance often declines on exome or panel data. SigMA [36] was developed specifically for panel sequencing and moves HRD prediction closer to clinical use, but it focuses primarily on single base substitution (SBS) signals and omits structural alterations, despite evidence that some of this information can be retrieved from targeted panels [37,38]. Overall, many existing tools either depend on input features that are difficult to obtain in standard diagnostics or are evaluated in settings that do not reflect routine clinical sequencing, limiting their practical utility.

Here, we present a systematic comparison of end-to-end DL and feature-based ML for genomic biomarker prediction using data modalities that are realistically available in clinical practice. We implement supervised and self-supervised DL encoders for small somatic variants and gene-level CNVs and benchmark them against feature-based ML models [18,32,39]. We evaluate MSI prediction from unfiltered somatic mutations in colorectal (COAD, READ), gastric (STAD), and uterine cancers (UC), and HRD prediction from combined mutation and CNV profiles in breast (BRCA), ovarian (OV), prostate (PRAD), lung (LUAD, LUSC), endometrial (EC), and pancreatic (PAAD) cancers. To reflect clinical sequencing practice, we assess performance not only on whole-exome sequencing (WES) but also under pseudo-panel mutation subsets and on real targeted-panel data. Finally, we use explainability methods to characterize the genomic features that drive predictions and to relate them to established MSI- and HRD-associated mutational patterns.

Together, this work benchmarks DL and classical ML across exome and targeted-panel sequencing, provides practical guidance for model selection and evaluation, and assesses interpretability. Overall, our findings support the use of these models as complementary tools in translational clinical genomics, enabling the extraction of additional value from routinely generated sequencing data.

## Methods

### Data acquisition

SNVs and small indels of 4,925 WES cancer patients from The Cancer Genome Atlas Program (TCGA) (n=4,440 patients) and the Clinical Proteomic Tumor Analysis Consortium (CPTAC) (n=485 patients) were utilized. (**Fig. 1a**) TCGA controlled-access and CPTAC public mutation data were obtained in Mutation Annotation Format (.maf) files through the Genomic Data Commons (GDC). Panel data was acquired for GENIE (n=221) version 18.0 [40] and two GeneseeqPrime® cohorts (C2 n=85 & C3 n=416) [41]. In total we acquired 7,554,787 million somatic SNVs and small indels for TCGA, 973,661 mutations for CPTAC and 5897 mutations from GENIE. CNV information for HRD experiments was captured in the form of WGS and panel segment files (**Fig. 1b**).

**Fig. 1.**
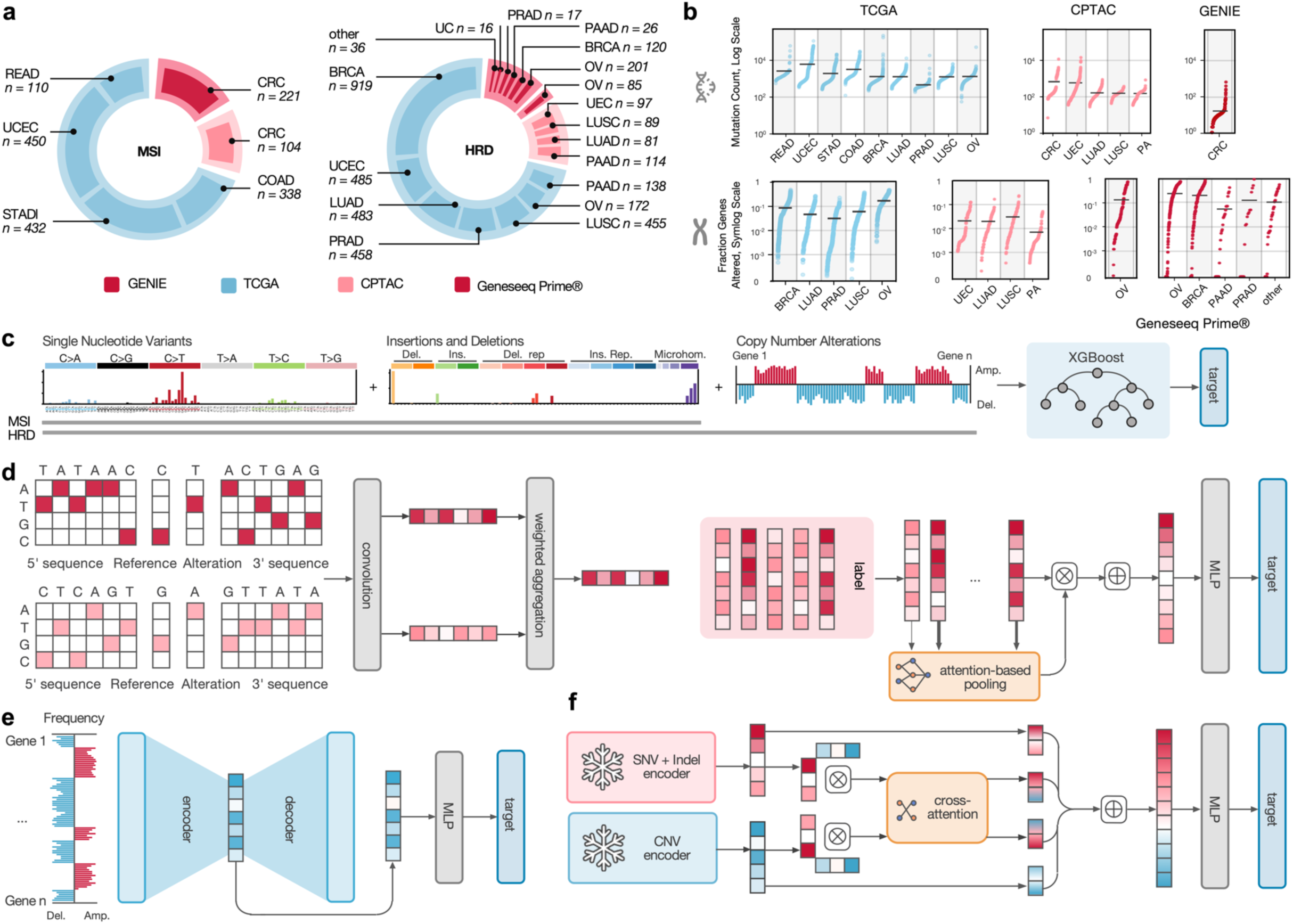
Overview of the study setup. **a** Datasets (TCGA and CPTAC) used in this study with cancer types and patient counts stratified by biomarkers. Cancer types from TCGA are accentuated in blue, red indicates CPTAC. **b** Somatic mutation counts and fraction of genes altered by CNVs by cohort and cancer type used in this study. The black line indicates the mean mutation count. **c** ML workflow. Aggregation of SBS and Indel features for MSI prediction. Addition of CNV features per gene for HRD predictions. The ML model of choice is XGBoost. **d** AttMIL model. Mutations from (a) are separated into four blocks: upstream, reference, alteration and downstream element. Each block contains 20 nucleotides or gaps. The reverse strand is modeled in the same manner by the reverse complement of the mutation. Both strands are passed through a 2D-convolution and a dense layer producing a mutation feature vector. Mutation vectors are gathered at the patient level and are aggregated by the attMIL mechanism to a patient feature vector. Patient features are used for the classification task. **e** CNV autoencoder. CNVs in the form of gene level amplifications/deletions are compressed into a patient level feature vector by an autoencoder. These representations are used for the classification task. (COAD - colon adenocarcinoma, READ - rectal adenocarcinoma, STAD - stomach adenocarcinoma, UCEC - uterine corpus endometrial carcinoma, OV - Ovarian serous cystadenocarcinoma, BRCA - Breast invasive carcinoma, PRAD - Prostate adenocarcinoma, PAAD - Pancreatic adenocarcinoma, MLP - multilayer perceptron, CRC - colorectal cancer, PA - pancreatic adenocarcinoma, UCE - uterine endometrial carcinoma, SNV - single nucleotide variant)

To reconstruct the mutation sequence context, mutation entries were mapped back to their genomic locations using the ‘Chromosome’, ‘Start_Position’, and ‘End_Position’ columns to generate their sequence context. Human reference genome builds GRCh37 was used for TCGA and GENIE, and GRCh38 for CPTAC data respectively, with chromosome sequences obtained from the Ensembl (https://www.ensembl.org/) genome database. CNV segments from WGS were filtered regarding exonic regions with the help of Gencode genome annotation release 10/21 (https://www.gencodegenes.org/) to imitate WES based CNVs.

For prediction of MSI status, four common cancer types were selected: COAD (n=338), READ (n=110), STAD (n=432), and UCEC (n=450) (**Fig. 1a & S1**), resulting in a dataset of 1,330 patients from TCGA. MSI status was obtained through cBioPortal (https://www.cbioportal.org/) and previous studies conducted on TCGA data [42]. The MSI status of all patients was determined using consensus calls from polymerase chain reaction (PCR) -based assays, MANTIS [43], and MSIsensor [44], with previously described thresholds set at binarized values of 0.4 and 3.5, respectively (**Additional File 1, Table S1**). For validation, 104 COAD patients from CPTAC were used (**Table S2, Additional File 1**). To assess model performance on more challenging cases, the trained models were also applied to the TCGA non-consensus patients (n=46). Furthermore, we utilized GENIE CRC (n=221) as another dataset. (**Table S3**)

For prediction of HRD status, seven TCGA cancer types where HRD is found were selected [34]: BRCA (n=919), UCEC (n=485), LUAD (n=483), PRAD (n=458), LUSC (n=455), OV (n=172) and PAAD (n=138) (**Fig. 1a, Fig. S2**). The genomic HRD scar score [45] consists of three numerical properties: loss of heterozygosity (LOH) [46], telomeric allelic imbalance (TAI) [47], and large-scale transitions (LST) [48] from which a sum HRD status was determined. Using the general cutoff of 42 [49], we generated binarized prediction targets (**Additional File 2, Table S4**). CPTAC was used as an external validation cohort. Here we applied our models to PAAD (n=114), LUAD (n = 81), LUSC (n=89) and UCEC (n=97) (**Table S5**). To further test capabilities in panel data, we utilized a dataset of two cohorts analyzed with GeneseeqPrime® HRD, with cohort C2 consisting of only OV cancers (n=85) and C3 of mixed cancer types (OV n=201, BRCA n=120, PAAD n=26, PRAD n=17, UC n=16, other n=36) (**Table S6&7**) [41].

### Data Preparation

For the ML models, SNVs were represented in their SBS and indel forms, comparable to Catalogue Of Somatic Mutations In Cancer (COSMIC) v3.4 [50] features. Therefore, we used 179 input features for the ML model, 96 for SBS mutations and 83 for indels. Feature values were normalized by mutation count of the respective patient. Gene-level CNVs were obtained by averaging the log₂-transformed copy-number ratios of all segments per gene resulting in 19,444 features ordered linearly by appearance on the chromosomes from chromosome one to Y (**Fig. 1c**). Because GeneseeqPrime® reports gene-level copy-number as discrete copy states (e.g., loss/neutral/gain), we mapped TCGA/CPTAC CNVs to the same representation and retrained the CNV classifier before evaluating on GeneseeqPrime®. (**Supplementary Methods**)

The input data for the small somatic mutation encoder consisted of the sequence context surrounding each mutation. A 20-nt window was extracted around each mutation for both the mutated and reference alleles, along with 20-nt upstream and downstream context. Shorter mutations (e.g., SNVs) or reference sequences (in case of insertions) were padded with ‘–’ to match the window size. For all experiments, a window size of 20 nucleotides was defined to capture the specific indel signatures associated with MSI and HRD mechanisms (**Fig. 1d**). To account for the double stranded nature of DNA, the reverse complement of each mutation sequence was provided as well. All sequences were numerically encoded by mapping nucleotide bases (A, C, G, T) and gap characters (’-’).

For the CNV encoders we tried two approaches. A MIL approach in which CNV segments are the input, and different self-supervised models on gene level CNVs. For the former, each instance was made up of seven features: the log₂-transformed copy-number ratios, segment center position ((end position - start position) / 2), segment length, segment position relative to the chromosome (chromosome length / center position), chromosome, is_amplification and is_deletion (cutoff < -0.2 and > 0.2). For the latter, CNV data was represented as for the ML case in the form of gene-level feature vector per patient. (**Supplementary Methods**)

Furthermore, to simulate how our models perform on mutations from targeted sequencing panels, we filtered the WES-derived variant data using the gene lists from FoundationOne CDx (FO) (324 genes) [40,41] and TruSight Oncology 500 (TSO) (523 genes) [42,43]. Variants were filtered by matching their HUGO nomenclature entries to the gene lists. Since the gene panels both include only coding exonic regions, we kept mutations in the TCGA data with the following variant classification: Missense_Mutation, Silent, Nonsense_Mutation, Frame_Shift_Del, Frame_Shift_Ins, In_Frame_Del, In_Frame_Ins. In CPTAC CRC, after filtering for TruSight Oncology 62,860 variants from 432 genes remained and after the FoundationOne 39,960 variants from 280 genes remained. Because the attention-based MIL model was trained under a characteristic range of bag sizes, strongly reduced instance counts represent an additional out-of-distribution shift that can confound comparisons. To control for this bag-size effect, we used a simple upsampling procedure in which the remaining mutation instances were repeated to match a target instance count range observed in the training data. Therefore, based on the % of exomes covered by the sequencing panels, TSO data was upsampled 30 times and FO data 50 times. Since GENIE includes a mixture of panels, we used a 10x factor as default.

### Data Split

To enhance generalizability and mitigate potential site-specific biases, a site-specific training split based on the Tissue Source Site (TSS) codes within the TCGA patient identifiers was conducted [51]. TSS codes were mapped to their respective institutions (**Additional File 3**). The test set for each model comprised patients from specific institutions excluded from the training set to ensure comparable results. The training data was divided for a 5-fold cross-validation in an approximate 70:15:15 split (**Fig. S1a & S2a**). In the training dataset, positive samples for MSI ranged from 20 to 22 % between folds, and from 15 to 17 % for HRD (**Fig. S1b & S2b**).

### Baseline Machine Learning Classifiers

To establish ML a classifier on SNVs as baseline for performance, we compared logistic regression (LR), support vector machines (SVM), random forests (RF) and extreme gradient boosting (XGBoost). The parameters set in the scikit-learn v1.2.0 and xgboost v2.1.1 packages in Python were: LR - LogisticRegression(max_iter=2000, C=1.0, penalty=“l2”), SVM - LinearSVC(C=1.0), RF - RandomForestClassifier(n_estimators=500, max_depth=None, min_samples_leaf=1,n_jobs=-1, random_state=42), XGBoost - XGBClassifier(n_estimators = 100, use_label_encoder = False, eval_metric = logloss, random_state = 42, learning_rate = 0.01, max_depth = 6, subsample = 0.8, colsample_bytree = 0.5) models were employed. We applied the models to the HRD predictions as this is the harder task and a differentiation between the models is better visible. Final predictions in the single modality and multi-modality case were conducted on the best performing model of the previous ablation, the XGBoost. In the multi-modal case we concatenated the 179 SNV and 19444 CNV features. (**Fig. 1c, Table S8**)

### Deep Learning Setup

The small somatic mutation encoder followed the implementation by Anaya et al. [32] in TensorFlow v2.12.0 and consisted of a trainable mutation encoder coupled with an attention-based multiple-instance learning (attMIL) module (**Fig. 1d**) (**Supplementary Methods**). The mutation encoder applies a 2D convolution to the forward and reverse-complement sequence windows separately. The resulting representations are concatenated and passed through a dense layer, yielding a 128-dimensional embedding for each mutation. In the attMIL module [52], all mutation embeddings of a patient, in the form of ragged tensors, are aggregated into a single patient-level vector using an attention mechanism with weighted-sum pooling. The resulting 128-dimensional patient embedding is fed into a multilayer perceptron (MLP) for classification. For MSI models, training was conducted for up to 300 epochs with a learning rate (LR) of 1e-3. HRD models were trained for up to 1000 epochs with a LR of 1e-4. (**Fig. 1d**)

We applied a similar attMIL scheme to segment-level CNV data. Each of the seven segment features (log₂ ratio, segment center, segment length, chromosomal position, chromosome index, amplification flag, deletion flag) was first linearly embedded. Chromosome features were embedded into eight dimensions; all other features into one dimension. The concatenated instance embedding was then aggregated with attention into a patient-level representation for downstream classification. (**Supplementary Material**)

Gene-level CNVs were encoded using self-supervised models, grouped into autoencoders (AEs) and masked-language modeling (MLM) approaches. For autoencoders, the 19,444-dimensional gene vector was compressed through three dense layers into a 512-dimensional embedding. Smaller embeddings led to information loss, whereas larger ones increased overfitting. AEs were trained fold-wise (LR = 1e-3, max. 200 epochs). Supplementing training with CNV profiles from additional TCGA samples did not improve performance. Patient embeddings produced by the AEs were fed into a two-layer MLP for classification. (**Fig. 1e**) Variational AEs and masked autoencoders (masking rate 30%) were also evaluated. (**Table S9**)

For MLM, we adapted the BulkRNABert architecture [18] to CNVs by binning log₂ CNV values into 64 discretized bins (vocabulary size 67 including CLS, MASK, PAD) and adding a gene embedding. Fifteen percent of input positions were randomly masked. A BERT-style transformer [53] was trained to reconstruct the masked bins (LR of 2e-4, max. 50 epochs). Patient-level embeddings were generated by taking the mean of the token embeddings. We also evaluated a Hyena-based encoder [54], replacing transformer attention with convolutional mixing. Both MLM models performed best with 512-dimensional embeddings, matching the AE embedding size for comparability. (**Table S9**) (**Supplementary Material**)

For multimodal prediction, we combined the pretrained SNV attMIL encoder with the best-performing CNV encoder (the simple AE). Both encoders were frozen and fused using a cross-attention layer. (**Fig. 1f**) Alternative strategies, such as simple concatenation or joint finetuning with warmup (1e-7 to 1e-5), performed worse than frozen encoders with cross-attention. (**Table S10**)

Binary classification used a sigmoid output with binary cross-entropy loss and the Adam optimizer. All models were trained on an NVIDIA RTX A6000 GPU (46 GB) and A100 SXM4 (80GB).

### Statistical Analysis

Model performance was evaluated using accuracy, F1 score, receiver operating characteristic (ROC) area under the curve (AUC), and precision-recall (PR) AUC, specificity and sensitivity for classification tasks. To convert continuous prediction scores into binary labels, we selected a decision threshold on the training set by maximizing Youden’s *J* statistic on the ROC curve (i.e., maximizing sensitivity + specificity − 1) (**Formula 1**). The threshold was determined within each training fold and applied to the corresponding held-out fold and external data.

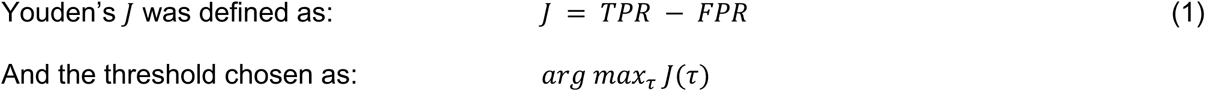

Where *TPR* is the true positive rate (sensitivity) and *FPR* is the false positive rate (1- specificity).

To compare model performance, we primarily used the F1 score and PR-AUC, as they provide more reliable estimates in imbalanced datasets than accuracy or ROC-AUC and are less affected by threshold selection than sensitivity or specificity.

To assess whether the best-performing model significantly outperformed the others, we carried out paired comparisons across cross-validation folds using the Wilcoxon signed-rank test on the F1 scores. All comparisons were evaluated between identical test sets, ensuring paired observations. Reported p-values correspond to two-sided tests. Because 5-fold cross-validation yields only five paired performance values, classical significance tests have limited statistical power. For the Wilcoxon signed-rank test, the smallest attainable non-zero p-value with n=5 is 0.0625, meaning that nominal thresholds such as p < 0.05 cannot be reached even in cases of large and consistent effect sizes.

### Explainability Techniques

Several explainability techniques were applied to interpret the models’ predictions. This approach allowed us to interpret how models encode and evaluate SNVs and CNVs, as well as how these encodings contribute to patient-level predictions.

For XGBoost models, Shapley additive explanations (SHAP) values were calculated with the python library shap (v0.45.1) for each input feature. By default, 20 most important features are ranked by SHAP value and displayed in increasing order.

For the SNV DL encodings, dimensionality reduction was performed using Uniform Manifold Approximation and Projection (UMAP) (python library umap-learn v0.5.5), which facilitated the visualization of mutation- and patient-level feature vectors by projecting them into lower-dimensional space to identify sample clusters. Mutation-level features were extracted from a layer that encodes individual mutations into vector representations, while patient-level features were obtained from the aggregation layer, which integrates information across all mutations for a given patient. Mutation encodings were then stratified based on attention value and grouped using k-means clustering (k=7) with the sklearn.cluster library, allowing the analysis of the importance of similar mutations to the model. The choice of k=7 balanced sufficient granularity to capture distinct mutation patterns with avoiding excessive fragmentation. To further understand the relationship between these clusters and biological processes, mutation catalogs from the clusters were generated and compared to known COSMIC mutational signatures associated with dMMR and HRD. Additionally, SigProfilerExtractor [55] was employed to extract SBS and Indel signatures from TCGA data, aiming to identify which signatures are present in the WES for better comparability, since COSMIC signatures were originally derived from WGS. Separate extractions were performed for the cancer types used in the MSI and HRD experiments. The parameters used were: exome=True, minimum_signatures=1, maximum_signatures=10, nmf_replicates=100, cpu=10.

To identify which gene-level CNV features contributed most to the AE’s reconstruction, we used Integrated Gradients (IG) [56] (**Formula 2**). IG attributes importance to each input feature by accumulating the gradients of the model’s output with respect to the input along a straight-line path from a baseline input *x*′ to the actual input *x*.

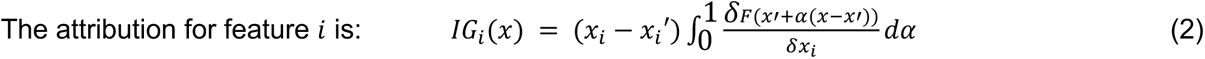

where *F* is the model, *x* is the sample, *x*′ is the baseline (zero CNV state).

In short, IG highlights genes where small changes in copy number strongly influence the model’s internal representation. To pick top influential genes, we set a threshold at the 99th percentile of importance. These genes were then evaluated by chromosomal position and function to understand if the model bases its predictions on certain genomic properties.

Finally, the relationship between the model’s prediction scores and various tumor properties was examined for potential trends. For MSI, these included associations with cancer type, tumor mutational burden (TMB), indel mutation density, driver mutations in dMMR genes (e.g., *MLH1*, *MSH2*, *MSH3, MSH6*, *PMS2*) and *POLE*, as well as the methylation status of *MLH1* promoter. For this, TCGA patients included in the model’s validation and test sets were selected. For HRD, correlations were evaluated in relation to cancer type, *BRCA1/2* status, continuous genomic scarHRD scores and their components (i.e., LOH, TAI, and LST), TMB, CNV burden in known hotspots (3p/5q/8p loss, 3q/8q/17q/20q gain) [57] and mutations in additional HR pathway genes (e.g., *BRCA1, BRCA2, ATM, CHEK2, PALB2, RAD51C*).

### Score calibration for panel evaluation

Platt calibration was used for HRD in cross-assay panel evaluation. After training the base prediction models on TCGA, we fitted a logistic regression calibration model (Platt scaling) on the C3 GeneseeqPrime® multi-entity to map raw prediction scores to calibrated probabilities C2 GeneseeqPrime® ovarian dataset [58]. We did not perform calibration for MSI because GENIE and splitting it for calibration risked overstating performance. Calibration quality was assessed using calibration curves and the Brier score, reported before and after calibration.

## Results

### MSI predictions from unfiltered somatic mutations

MSI arises because of defects in the mismatch repair (MMR) pathway [59,60], leading to characteristic small-scale indels in repetitive sequences and distinctive SBS patterns (**Fig. 2a**), which in principle are recognizable by DL-based models [9,61–63]. We trained an attMIL model to detect MSI status in 1,330 TCGA patients (COAD, READ, STAD and UCEC), and then externally validated it on the CPTAC COAD cohort (n=105). The attMIL model achieved an F1 score of 0.97±0.01, a PR AUC of 1.00±0.00, sensitivity of 0.95±0.02, and a specificity of 1.00±0.00, confirming its ability to detect MSI-related patterns also on external datasets (**Table S11**).

**Fig. 2.**
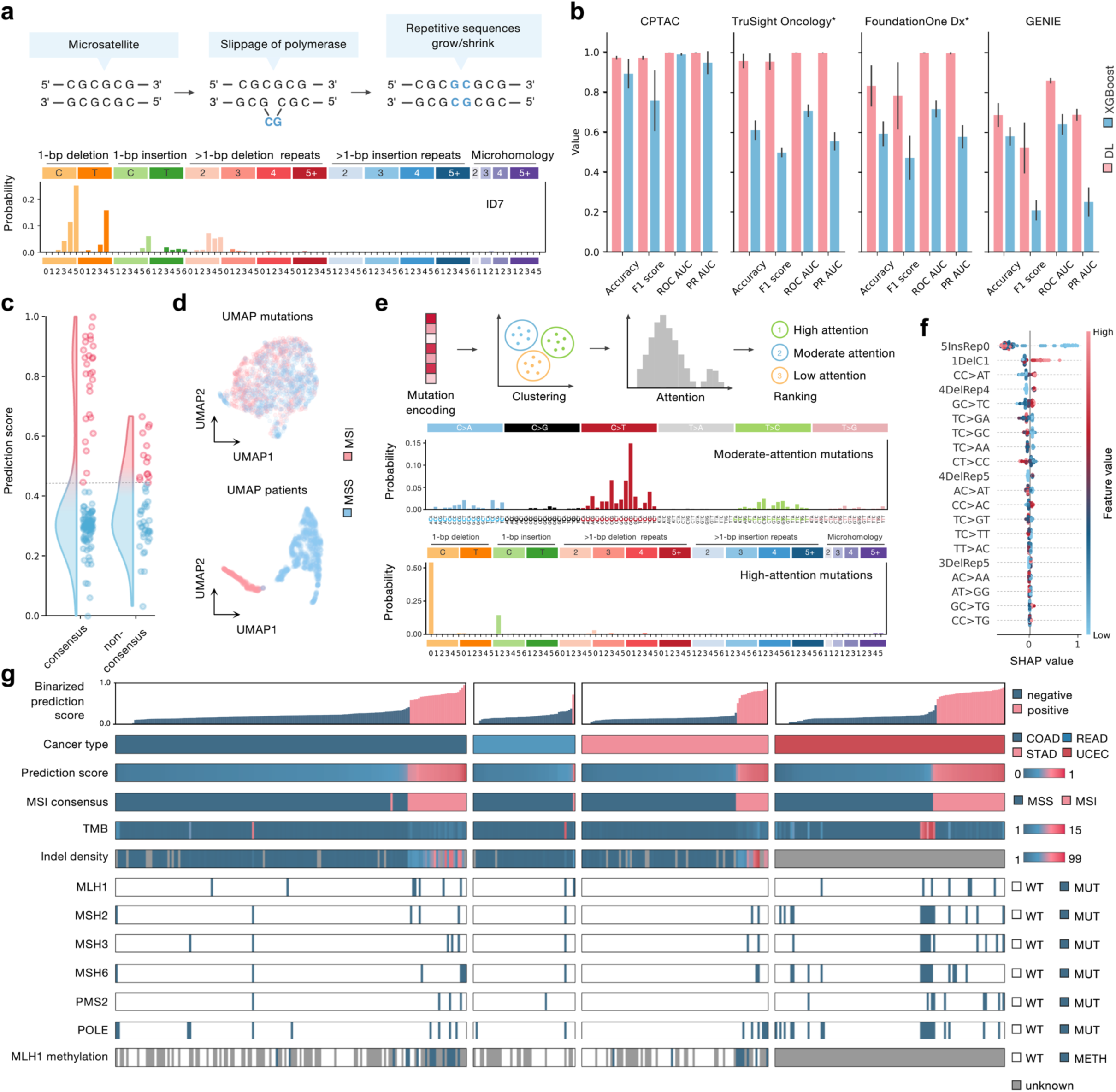
DL predicts MSI with high accuracy in panel sequencing. **a** Mutation mechanism and COSMIC Indel signature of MSI. **b** Bar chart of performance metrics to compare attMIL and XGBoost models on the full external CPTAC dataset, CPTAC data filtered by targeted sequencing panels of FoundationOne and TruSight Oncology (* indicates pseudo-panels) and GENIE CRC panel data **c** DL model prediction scores of the attMIL model comparing consensus and non-consensus labeled cases between PCR status, MSISensor and MANTIS score. **d** UMAP of mutation and patient level features. **e** Explainability of the attMIL model. 5,000 random mutation features were clustered by k-means (k=7) and clusters were ranked by mean attention score. SBS/Indel mutation catalogues of two mutation clusters. **f** Shap values of the XGBoost models’ 20 most important features in descending order. **g** Association of attMIL model prediction scores with genomic properties of the patients separated by cancer type. Comparisons are made between the prediction scores, the ground truth, mutation densities (general and indel specific), amino acid changing mutations and methylations in MMR associated genes. (COAD - colon adenocarcinoma, READ - rectal adenocarcinoma, STAD - stomach adenocarcinoma, UCEC - uterine corpus endometrial carcinoma, MSI - Microsatellite instability, MSS - microsatellite stability, WT - wildtype, MUT - mutated, METH - methylated)

Next, we compared the attMIL performance with that of a SOTA ML model (XGBoost) to investigate whether DL is indeed more powerful than standard ML for this application. We chose XGBoost as a reference since it performed best among our group of classical ML models (**Table S8**). While XGBoost also performed well on the CPTAC dataset (F1 score of 0.76±0.15, PR AUC of 0.95±0.06, sensitivity of 0.99±0.02, specificity of 0.80±0.16; **Fig. 2b**), a difference in performance between DL and ML models was observed when comparing F1 scores with a p-value of 0.06, the minimum possible p-value for our Wilcoxon signed-rank test. Thus, although MSI prediction is relatively straightforward for classical ML, the DL approach is still substantially more accurate. (**Table S11**)

Given the similarities in high-level performance, we further investigated the models’ performance in more challenging MSI cases. Thus, we applied both models to previously excluded non-consensus patients (n=46), where PCR, MSIsensor, and MANTIS results were discordant. As expected, the DL model’s prediction scores in these cases clustered closer to the classification threshold, indicating lower confidence (**Fig. 2c**). A similar trend was observed with XGBoost, where several patient’s scores also grouped towards the threshold (**Fig. S3a**). Notably, the attMIL model identified a similar amount of MSI cases as XGBoost (7 and 5 from 46 patients). However, Venn diagrams comparing model outputs with MSI detection methods revealed minimal overlap, highlighting the ambiguity of these non-consensus cases and the need for further validation (**Fig. S3b**).

### MSI is detectable from sequencing panels

In the real world, most patients do not undergo WES, but panel sequencing of a few hundred genes. We first evaluated the performance of DL and ML models on pseudo panel sequencing data. To this end, the number of somatic mutations was filtered to include only genes present in two targeted sequencing panels: FoundationOne Dx [64,65] and TruSight Oncology [66,67]. Reducing the mutation set to the TruSight Oncology 523 gene panel revealed a significant divergence in performance between DL and ML models, with the attMIL model achieving an F1 score of 0.96±0.04 outperforming XGBoost’s 0.50±0.02 (p-value 0.06) and a PR AUC of attMIL of 1.00±0.00 compared to 0.56±0.05 of XGBoost. As expected, the task became more challenging with fewer mutations available, indicated by lower overall F1 score - however, the DL model maintained a reasonably good performance. When the genes were further narrowed to 324 from the FoundationOne Dx panel, performance differences persisted despite increased variance (F1: attMIL 0.78±0.17 vs. XGBoost 0.47±0.11, p-value 0.13; PR AUC: attMIL 1.00±0.00 vs. XGBoost 0.58±0.06). (**Fig. 2b, Table S11**)

To assess whether these findings extend to real-world panel data, we applied both models to the GENIE CRC cohort. The DL model again outperformed XGBoost (F1: attMIL 0.52 ± 0.13 vs. XGBoost 0.21 ± 0.05, p-value 0.06; PR AUC: attMIL 0.68 ± 0.03 vs. XGBoost 0.25 ± 0.07), suggesting better generalization of the DL approach to this panel setting. As expected, overall performance was lower than in the WES-based experiments, consistent with differences in sequencing protocols and mutation calling, and with the impact of very low mutation counts in some samples. When restricting the analysis to patients with at least 10 mutations, performance partially recovered (F1 0.60 ± 0.14, PR AUC 0.84 ± 0.03). Together with the pseudo-panel results, these findings indicate that the DL model maintains an advantage over classical ML for MSI prediction in clinically relevant genomic subsets present in commercial sequencing panels. (**Fig. 2b**, **Table S12**)

### DL and ML detect biomedically relevant MSI patterns

Subsequently, we investigated if the feature representations obtained by the DL model result in a clustering of patients, which would provide further proof that the DL model learned clinically relevant patterns. We investigated a possible clustering of at both the mutation and patient levels (**Fig. 2d**). While no substantial differences were apparent at the mutation level, a clear separation emerged at the patient level. MSI and MSS cancers segregated distinctively, indicating that the DL model was able to encode their respective patients differently. Furthermore, the patient level features exhibited tissue specificity forming distinct clusters in UMAP space (**Fig. S3c**). Despite being trained only on mutation data, the model inherently captured patterns associated with tissue types.

We then evaluated which mutations were most influential for the model’s predictions by extracting mutation encodings from an embedding layer. These encodings were clustered using k-means (k=7) and ranked by the mean attention values of their mutations to identify highly and minimally influential groups (**Fig. 2e**). When examining the mutation catalogs of the clusters with the highest attention scores, they were enriched in C and G indels. These mutations were also captured as mutation signatures by SigprofilerExtractor and could be related to the COSMIC indel mutation signature ID7 [50] (**Fig. 2a**), related with MSI, which depicts deletions of mononucleotide stretches but also of dinucleotide repeats. Altogether, this suggests that among patient mutation profiles, the model captures relevant patterns, associated with MSI.

When analyzing a mutation cluster with average attention scores, we observed several similarities to the SBS6 and SBS15 mutation signatures associated with defective MMR (dMMR) [50] (**Fig. S4a**), also found in the WXS de-novo data by SigProfilerExtractor (**Fig. S4b**). Similarities between the mutation cluster and signatures encompassed the features of the C>T mutation class, with comparable peaks in the sequence context pairs of an up- and downstream GG, AG and CG. This could further indicate that the model may have learned to group and assign weights to MSI-specific mutations. However, the model’s focus is laying more on indel mutations than SNVs. Comparing this to the Shapley additive explanation (SHAP) analysis of the features of XGBoost, XGBoost also utilizes deletions of a single C/G similar to the DL model (**Fig. 2f**). However, other features that strongly influenced the XGBoost model’s predictions, such as indels at repetitive sites or certain C>G/C>A mutations, did not resemble classical dMMR signatures, suggesting either the identification of novel patterns or a potential bias towards irrelevant features.

### DL identifies MSI-associated patterns beyond mutation counts

Finally, we investigated whether the model’s predictions were driven by subtle patterns within the mutations or if it was simply confounded by factors such as mutation counts. To this end, we investigated if the attMIL model’s predictions are associated with other relevant properties of the patients’ cancers related to MSI (**Fig. 2g**). These properties included the ground truth label, features of genomic instability, driver mutations, and the methylation status of MMR-related genes. The model’s normalized prediction scores were highly indicative of MSI and MSS status across all cancer types, effectively reflecting the underlying distinction between the two groups. Next, as expected, a correlation trend between the prediction score and TMB and indel density was observed. However, patients with high TMB/indel count in an MSS context were not predicted as MSI by the DL model. This suggests that the model learns subtle, clinically relevant patterns and does not simply count mutations as indicators for MSI. We also observed that driver mutations in key MMR pathway proteins are not necessarily indicative of a positive model prediction, nor were they always correlated with consensus labels from MANTIS, MSIsensor or PCR (**Fig. 2g**). A close relationship was found between the model scores and the MLH1 promoter methylation, as silencing of this gene has a more significant impact on MMR function than mutations alone [68].

These results emphasize DL’s potential as an alternative tool for analyzing NGS data, specifically for panel sequencing data. Our results demonstrate high performance, generalizability, and explainability of the DL-based attMIL model.

### DL and ML predicts HRD and in multimodal manner

We next investigated HRD, which arises from defects in core homologous recombination (HR) genes and related repair pathways. When HR fails to resolve double-strand breaks, cells rely on alternative mechanisms such as microhomology-mediated end joining (MMEJ) and non-homologous end joining (NHEJ), leaving a mixture of small-scale mutations and larger structural alterations in the genome (**Fig. 3a**) [69,70].

**Fig. 3.**
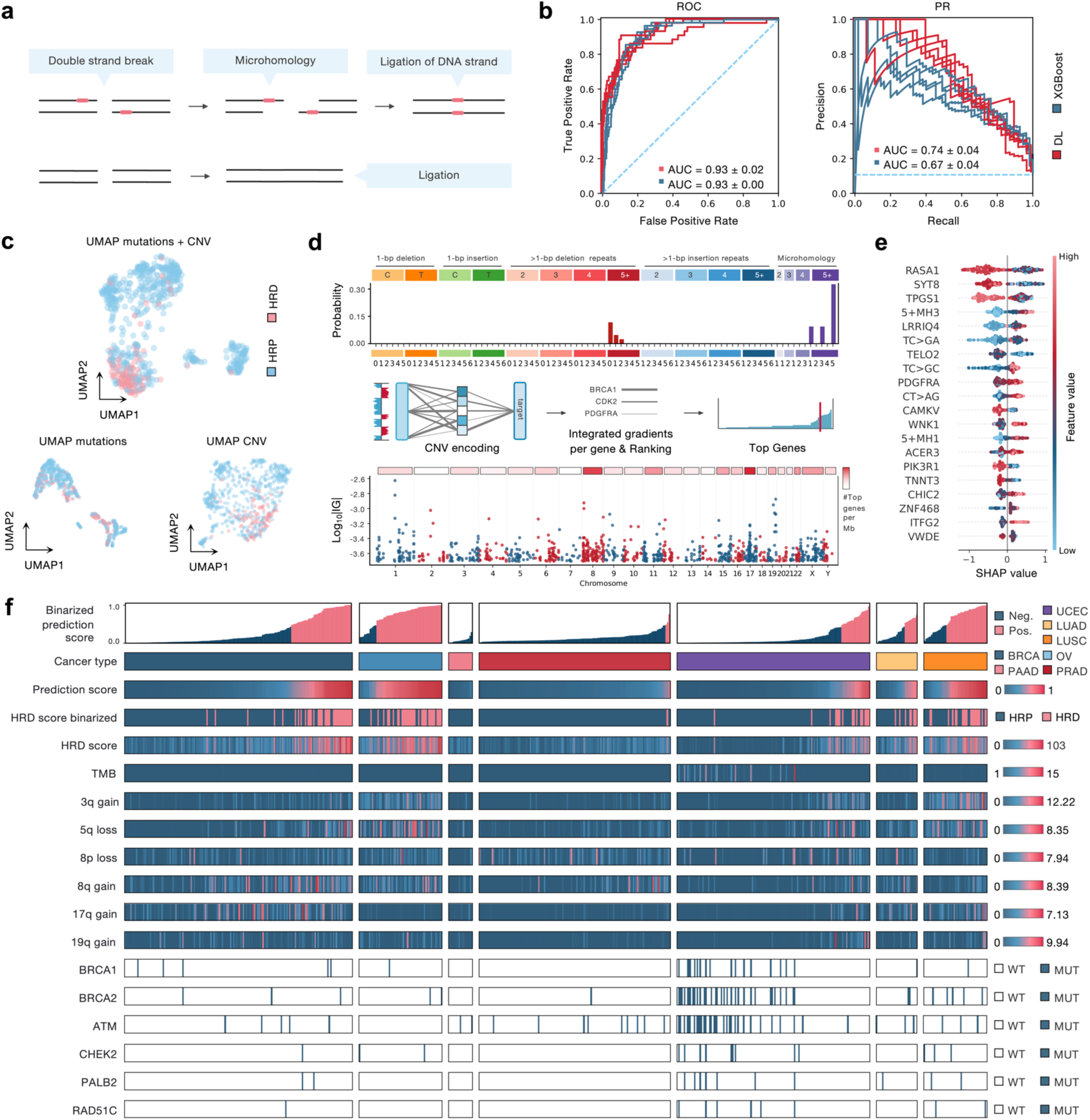
DL and ML predict HRD in a multimodal manner. **a** Mutation mechanism of MMEJ and NEHJ in response to HRD. **b** ROC AUC and PR AUC plots for attMIL and XGBoost HRD predictions. **c** UMAP patient level features of unimodal and multimodal models. **d** Shap values of the XGBoost models’ 20 most important features in descending order. **e** Indel mutation catalogues of the highest attention cluster and explainability of the autoencoder for CNVs. Integrated gradients were calculated for all weights in the autoencoder model. Gene level weights were calculated by the sum of all gradients for respective input. Gene level values were then ranked and top contributing genes were determined. **f** Association of multimodal DL model prediction scores with genomic properties of the patients separated by cancer type. Comparisons are made between the prediction scores, the ground truth, mutation densities (general and indel specific), CNVs in known chromosomal hotspots and amino acid changing mutations in HR associated genes. (OV - Ovarian serous cystadenocarcinoma, BRCA - Breast invasive carcinoma, PRAD - Prostate adenocarcinoma, PAAD - Pancreatic adenocarcinoma, HRD - Homologous Recombination Deficiency, HRP - Homologous Recombination Proficiency, WT - wildtype, MUT - mutated, IG - Integrated Gradients, CNV - Copy Number Variation)

We first trained DL and XGBoost models using only small somatic variants to predict HRD across BRCA, OV, PAAD, PRAD, LUAD, and LUSC in TCGA, with external validation on CPTAC UEC, LUAD, LUSC, and PAAD. On the external cohort, attMIL and XGBoost achieved comparable performance (attMIL F1: 0.41±0.04, XGBoost F1: 0.39±0.01, p=0.31; attMIL PR AUC: 0.35±0.03, XGBoost PR-AUC: 0.37±0.03) (**Table S14**). Performance was higher on the held-out TCGA test set (attMIL F1: 0.58±0.02, XGBoost F1: 0.58±0.02, p=0.44; attMIL PR AUC: 0.62±0.03, XGBoost PR AUC: 0.67±0.01; **Table S13**), illustrating that internal evaluation can substantially overestimate generalization for HRD and underscores the importance of external validation.

Because HRD is known to be strongly associated with structural changes, we next focused on copy-number information. Gene-level CNV models, implemented as an AE-based DL encoder and an XGBoost classifier, already outperformed the SNV-only models on the external validation cohort (AE DL F1: 0.56±0.05, XGBoost F1: 0.56±0.03, p=1.0; AE PR AUC: 0.67±0.03, XGBoost PR AUC: 0.69±0.08) (**Table S14**). However, DL did not provide an advantage over the ML baseline in this setting. Since TCGA CNVs were generated from WGS data and CPTACs CNVs as well, we expected good generalization.

We then combined SNV and CNV information in multimodal models. A cross-attention DL architecture using frozen pretrained SNV and CNV encoders was compared to a multimodal XGBoost model using both feature types. The multimodal DL model achieved the numerically highest performance (F1: 0.61±0.06, PR AUC: 0.74±0.04), while multimodal XGBoost reached F1: 0.58±0.02 and PR AUC: 0.67±0.04 (**Fig. 3b, Table S14**). The differences were not statistically significant (p=0.63), indicating that adding CNVs improves HRD prediction more than changing the model family itself. These experiments emphasize that for HRD, incorporating structural information is crucial, whereas DL and ML perform similarly once appropriate features are provided.

### Predictions highlight mutational patterns in alternative repair pathways

We next examined how the models represent HRD in feature space. UMAP embeddings of patient-level representations revealed a continuous gradient between HRD and HR-proficient (HRP) cases rather than a sharp separation (**Fig. 3c**). This gradient was most pronounced in the multimodal model and appeared less diffuse than in SNV-only or CNV-only embeddings. CNV-based embeddings showed a clearer HRD–HRP separation than SNV-based ones, consistent with the superior performance of CNVs for HRD prediction. Tissue-specific structure was again evident, with pronounced clusters for PRAD, PAAD, and OV, while BRCA spanned most of the UMAP space (**Fig. S5a**), indicating distinct mutational backgrounds across cancer types.

Attention scores from the SNV/indel attMIL encoder highlighted groups of mutations enriched for microhomology and deletions at 5-nucleotide repeats, consistent with COSMIC indel signatures ID6 and ID8 [50] (**Fig. 3d, Fig. S5b**) and a de novo indel signature recovered from our exome data (**Fig. S5c**). Both signatures are associated with alternative DSB repair pathways, including MMEJ and NHEJ, which are expected to be active in HRD tumors. Another group of highly attended mutations consisted mainly of diverse SBS classes related to SBS3 [50], in line with the elevated mutation burden characteristic of HRD cancers (**Fig. S5c,d**).

For the CNV encoder, we ranked genes by their integrated gradient attributions and mapped them back to genomic coordinates. This analysis revealed prominent hotspots on chromosomes 8 and 17, in agreement with reported chromosomal LOH on 17 and chromosomal instability on 8 in HRD-positive tumors [11] (**Fig. 3d**). Together, these findings show that the DL models base their HRD predictions on biologically plausible signatures in both small variants and CNVs.

The multimodal XGBoost model also made use of all mutation types. Among the top-ranked indel features were microhomology-related categories (e.g. 5+MH3, 5+MH1), which have previously been shown to be sufficient to classify HRD in isolation [63] (**Fig. 3e**). The model further emphasized certain C>G substitution features, consistent with HRDProfiler [35], and identified a diverse set of informative CNV genes, including *RASA1* (5q14.3), *SYT8* (11p15.5), *TPGS1* (19p13.3), *LRRIQ4* (3q26.2), *TELO2* (16p13.3), *CAMKV* (3p21.31), and *WNK1* (12p13.33). These genes are distributed across multiple chromosomal regions, suggesting that the model leverages a broad pattern of chromosomal instability rather than isolated events. Furthermore, several of these regions, including 3q and 5q, have previously been linked to HRD-based chromosomal instability [11,57,71].

Following the same approach as before with a MSI, we again compared the DL model’s predictions to target-specific tumor properties. (**Fig. 3f**). We observed that higher prediction scores correlated with a greater likelihood of patients being HRD-positive, suggesting that the model’s prediction scores reflect a ranking of HRD probability among samples. Upon examining the model’s prediction across cancer types, we noted that the model probably learned a tissue specificity as well (**Fig. S5a**). The model predictions were correlated to the general loss of 2q/3p/5q/8p gain of 3q/17q/20q but not as sole predictors leading to the conclusion that the DL models’ predictions are more fine grained than arm wise information [49,57]. Surprisingly, prediction scores and HRD scar scores were not closely correlated to mutations in BRCA1/2 or other pathway genes. UCEC even displayed a negative correlation with amino acid changing alterations in the pathway genes, highlighting the importance of HRD-caused pattern detection tools.

### CNVs as indicator for HRD status in panels

Compared to MSI, the overall lower performance for HRD is likely driven by the lower prevalence and more heterogeneous distribution of HRD-associated mutations across samples (**Fig. S6a**). In pseudo-panel experiments using only SNV features, performance approached random baseline, so we focused subsequent panel-like analyses on CNVs (**Fig. S6b**).

Gene-level CNVs in our study were derived from genome-wide CNV segments, whereas panel assays provide sparse, assay-specific coverage. CNV calling and segmentation from panels are therefore expected to be noisier and less precisely localized, introducing a domain shift relative to the TCGA CNV levels used for model development (**Fig. 4a**). Consistent with this, direct transfer of the TCGA-trained XGBoost CNV model to the GeneseeqPrime® ovarian cancer cohort performed poorly at the default decision threshold (F1: 0.42±0.27), despite a moderate PR AUC (0.83±0.06), suggesting that ranking information was partially retained but probabilities and thresholds did not transfer. After Platt calibration on the mixed entity GeneseeqPrime® cohort, XGBoost improved (F1: 0.62±0.10; PR AUC: 0.84±0.04), indicating that calibration can partially bridge the domain shift for this feature-based model (**Fig. 4b, Table S15**).

**Fig. 4.**
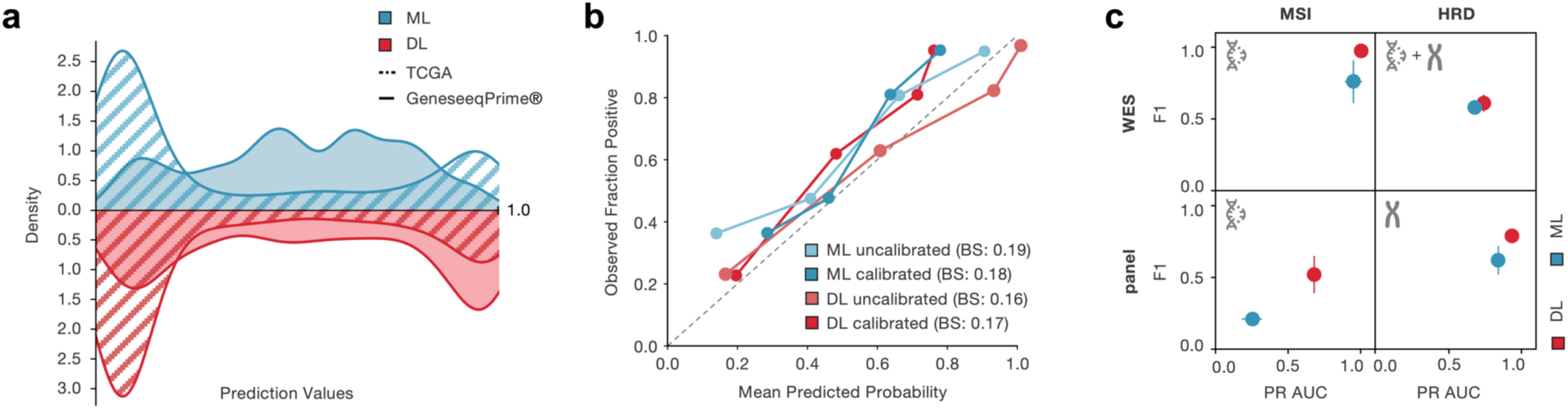
HRD predictions form panel CNVs. **a** Distribution of prediction values across ML and DL models and genome (TCGA) and panel (GeneseeqPrime®) data. **b** Calibration curves for un(calibrated) ML and DL models. Calibration was performed on the GeneseeqPrime® mixed cancer type cohort. Brier score was calculated for all calibration states as well. **b** Performance summary with F1 score and Precision-Recall AUC of experiments over model type, data modality and sequencing type. (BS - Brier score, PR AUC Precision-Recall area under the curve)

In contrast, the TCGA-trained autoencoder showed substantially better transfer to the panel cohort (F1: 0.75±0.09; PR AUC: 0.93±0.02) directly, underlined by a bimodal prediction value distribution, and calibration slightly improved its performance (F1: 0.78±0.04; PR AUC: 0.93±0.01) (**Fig. 4b, Table S15**). This suggests that the DL-CNV representations were more robust to assay-related distributional shifts than the feature-based XGBoost model. In practice, this implies that pretrained DL encoders may already provide useful HRD estimates in smaller in-house panel cohorts where full retraining is not feasible. When models were trained directly on the GeneseeqPrime® mixed cohort, both approaches achieved high accuracy (XGBoost F1: 0.90±0.02; AE F1: 0.93±0.01), confirming that CNV-based signal remains predictive on panels and that fine-tuning or retraining on panel-specific data remains preferable when sufficient panel samples are available (**Table S15**). In this setting, DL matched ML performance within the panel domain and transferred more robustly across assays, supporting its use for CNV-based HRD prediction in panel cohorts.

Taken together, these results show that MSI and HRD can be predicted from mutation-based profiles derived from genome, exome and clinical panel sequencing, but the optimal modelling strategy depends on the biomarker and data modality. MSI prediction consistently benefits from end-to-end DL, especially in panel settings, whereas HRD performance is primarily driven by the inclusion of CNV information. For HRD DL and feature-based ML perform similarly when trained within the same assay, while DL-based CNV encoders show greater robustness under cross-assay domain shifts. (**Fig. 4c**)

## Discussion

In this study, we used MSI and HRD as clinically relevant test cases to systematically compare end-to-end DL models and feature-based ML approaches for genomic biomarker prediction from tumor sequencing data across exome-wide and panel-like settings. DL models learned directly from somatic mutation and copy-number profiles without hand-crafted genomic features, and their internal representations aligned with biologically plausible patterns, including known indel and SBS signatures and HRD-associated CNV hotspots. For MSI, DL consistently outperformed ML across all settings, indicating that end-to-end sequence-based models are particularly effective for summarizing small-scale mutation patterns, especially in panel-like scenarios with sparse variant information. For HRD, the best performance was achieved by multimodal models that combined mutation and CNV data, where DL and ML reached similar accuracy when trained and evaluated within the same assay. However, when transferring models from exome-based data to panel cohorts, DL-based CNV encoders showed greater robustness than ML.

Together, these results provide practical guidance for selecting modeling strategies across biomarkers and sequencing settings. End-to-end DL represents a strong default when input data are sparse, when feature engineering is limited, or when models must generalize across assays. In contrast, well-tuned feature-based ML models can remain competitive when training and deployment occur within the same assay and sufficient data are available for retraining. By benchmarking against competitive ML baselines and validating performance on independent external cohorts, this work also highlights the importance of rigorous comparison and external validation when developing genomic AI methods for clinical use.

From a clinical perspective, these models are best viewed as complementary to, rather than replacements for, current diagnostic workflows. For MSI, immunohistochemistry remains a fast and inexpensive standard, but DL models operating on panel-based mutation data could provide additional evidence in ambiguous cases or when targeted sequencing is already available for other indications. For HRD, current diagnostic strategies typically combine sequencing of homologous recombination repair genes with genomic instability metrics, although both approaches have recognized limitations [72]. Given that HRD assessment already relies on sequencing data, models integrating small variants and CNVs from exome or panel datasets could provide an additional line of evidence without requiring further assays.

This study has limitations. Although we analyzed WES, WES-derived pseudo-panels, and clinical panel datasets, sequencing assays differ in design, coverage, and chemistry. These differences can shift the distribution of called mutations and CNV breakpoints and may affect model performance. For MSI, mutation upsampling in MIL for pseudo-panels could inflate performance on these artificial datasets. In HRD experiments, labels derived from composite genomic scar scores are imperfect and may introduce label noise and partial circularity when evaluating CNV-driven models. Future studies should therefore validate these models on additional panel cohorts, assess their association with treatment response, and explore integration with existing clinical pipelines.

Prospectively, our findings support the incorporation of AI models into clinical genomics workflows, where they can extract additional information from sequencing data that are already being generated for patients. Extensions to model architectures operating on raw sequencing reads, the development of genomic foundation models that provide reusable representations across tasks, and the integration of additional omics layers such as RNA expression or DNA methylation could further improve biomarker prediction. Ultimately, carefully benchmarked DL and ML models that run on routinely generated sequencing data may help make MSI and HRD assessment more consistent and scalable and may serve as a template for future NGS-based biomarkers in precision oncology.

## Conclusion

In summary, we present a systematic comparison of DL and ML approaches for genomic biomarker prediction across exome-wide and targeted-panel sequencing. Using MSI and HRD as representative test cases, DL models learned biologically meaningful patterns from mutation and copy-number data and matched or exceeded ML performance across most data regimes. Our results show that the benefits of DL are context-dependent and are most pronounced in settings with sparse inputs or cross-assay generalization. Together, these findings provide a practical framework for benchmarking, selecting, and evaluating AI models for genomic biomarkers beyond MSI and HRD in clinical and translational genomics.

## Abbreviations

AI: Artificial Intelligence
AUC: Area under the curve
AUROC: Area under the receiver operating characteristic
BRCA: Breast invasive carcinoma
COAD: Colon adenocarcinoma
CRC: Colorectal carcinoma
DL: Deep learning
HRD: Homologous Recombination Deficiency
HRP: Homologous Recombination Proficiency
LUAD: Lung adenocarcinoma
LUSC: Lung squamous cell carcinoma
MLP: Multilayer perceptron
MMEJ: Microhomology-mediated end joining
(d)MMR: (Defective) mismatch repair
MIL: Multiple instance learning
MSI: Microsatellite instability
MSS: Microsatellite stability
MUT: Mutated
NHEJ: Non-homologous end joining
NGS: Next-generation sequencing
PAAD: Pancreatic adenocarcinoma
PRAD: Prostate adenocarcinoma
OV: Ovarian serous cystadenocarcinoma
TCGA: The Cancer Genome Atlas
READ: Rectal adenocarcinoma
ROC: Receiver operating characteristic
UCEC: Uterine corpus endometrial carcinoma
WGS: Whole genome sequencing
WT: Wildtype
WXS: Whole exome sequencing
XGBoost: Extreme gradient boosting

## Declarations

## Ethics approval and consent to participate

This study was carried out in accordance with the Declaration of Helsinki. Datasets from CPTAC, TCGA, GENIE and GeneseeqPrime® do not require formal ethics approval for a retrospective study since samples are already anonymized. Furthermore, the analysis was approved by the Ethics commission of the Medical Faculty of the Technical University Dresden (BO-EK-444102022).

## Consent for publication

Not applicable.

## Availability of Data and Materials

The datasets supporting the conclusions of this article are available through the following sources: TCGA data is available at https://gdc.cancer.gov/about-data/publications/mc3-2017, and CPTAC data can be found at https://portal.gdc.cancer.gov/. TCGA access can be granted through eRA Commons (https://public.era.nih.gov/commonsplus/home.era?menu_itemPath=600) and dbGaP (https://www.ncbi.nlm.nih.gov/gap/). GENIE was requested through synapse.org https://www.synapse.org/Synapse:syn27056172/wiki/616601. CNV data of GeneseeqPrime® was shared as supplements in the respective publication [41]. Chromosome sequence information for data preparation is accessible at https://ftp.ensembl.org/pub/. Information regarding MSI and HRD status, as well as mutation statuses of driver genes, is available through cBioPortal at https://www.cbioportal.org/.

Scripts for data preparation and model implementation are hosted on GitHub at https://github.com/mxsunc/E2EDL-vs-ML-Biomarker-Detection. The code is written in Python and is platform independent (developed and tested on Linux-based systems). All required dependencies and version constraints are provided in the repository via `environment.yml`. The repository is released under the MIT License.

## Authors’ contributions

MU and JNK conceptualized the study. MU performed the DL experiments and explainability analysis. MU prepared the original draft and created the figures. JNK acquired funding and supervised the study. MU, CMLL, LZ, SS, TL, JV, AM, SF, TG, ZIC, JNK provided scientific input, reviewed and edited the manuscript.

### Acknowledgements

We acknowledge the TCGA (https://www.cancer.gov/tcga) and the CPTAC (https://proteomics.cancer.gov/programs/cptac) research networks, which generated the data on which the results shown in this manuscript are based on. The authors would like to acknowledge the American Association for Cancer Research and its financial and material support in the development of the AACR Project GENIE registry, as well as members of the consortium for their commitment to data sharing. Interpretations are the responsibility of study authors.

In accordance with the COPE (Committee on Publication Ethics) position statement of 13 February 2023 (https://publicationethics.org/cope-position-statements/ai-author), the authors hereby disclose the use of the following artificial intelligence models during the writing of this article: GPT-5.2 (OpenAI) for checking spelling and grammar.

JV acknowledges support from Fondation Inserm-Bettencourt and La Ligue contre le Cancer (these are my funders).

## Supplementary Material

## Supplementary Figures

**Supplementary Fig. 1.**
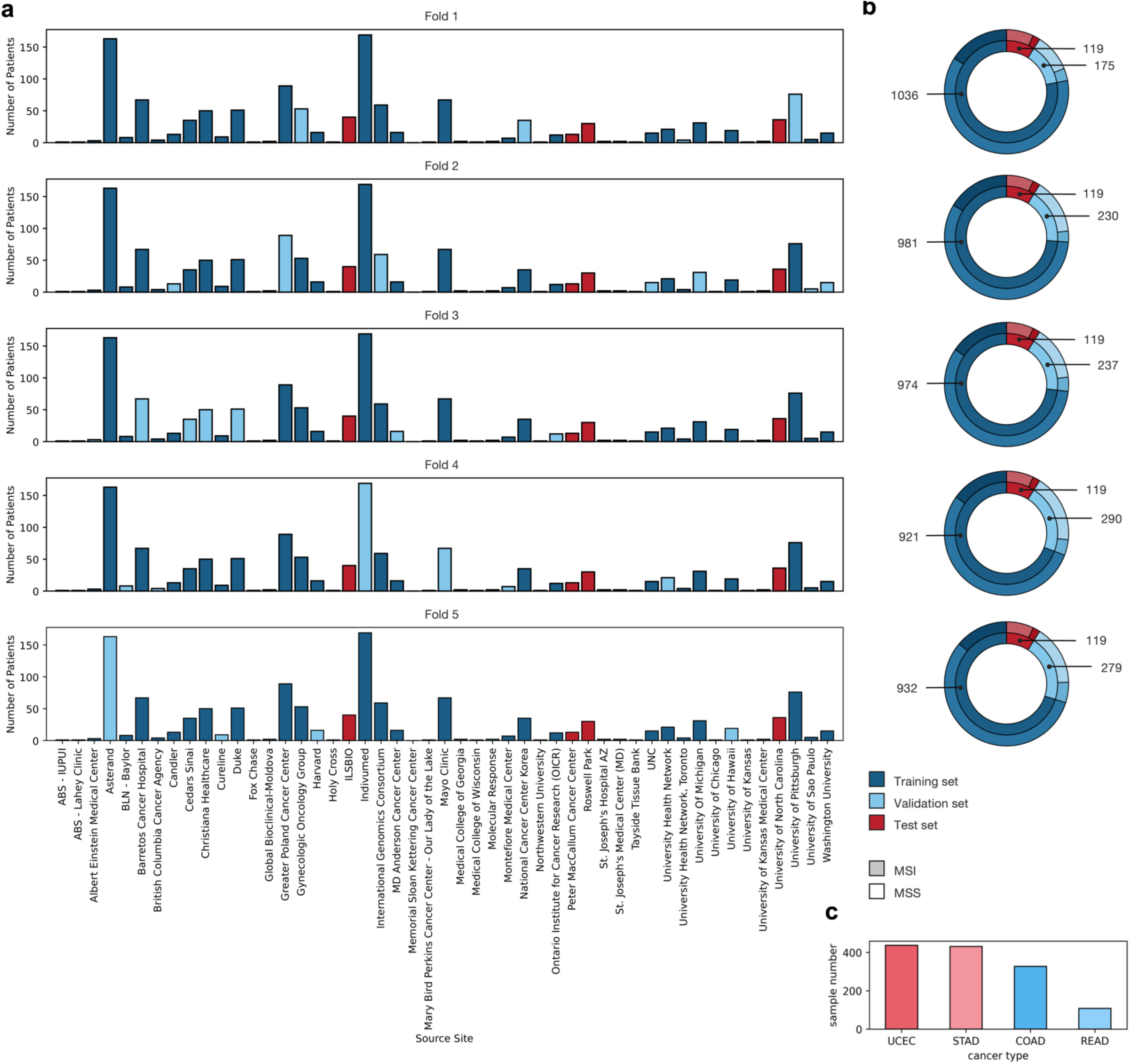
MSI data preparation. **a** Bar chart of site-specific data split for MSI. Dark blue indicates training data, light blue validation data and red the test set. The test set stays the same over all folds. **b** Doughnut plot of the data splits regarding class distribution of MSI vs MSS and number of patients. **c** Bar chart of number of patients per cancer type.

**Supplementary Fig. 2.**
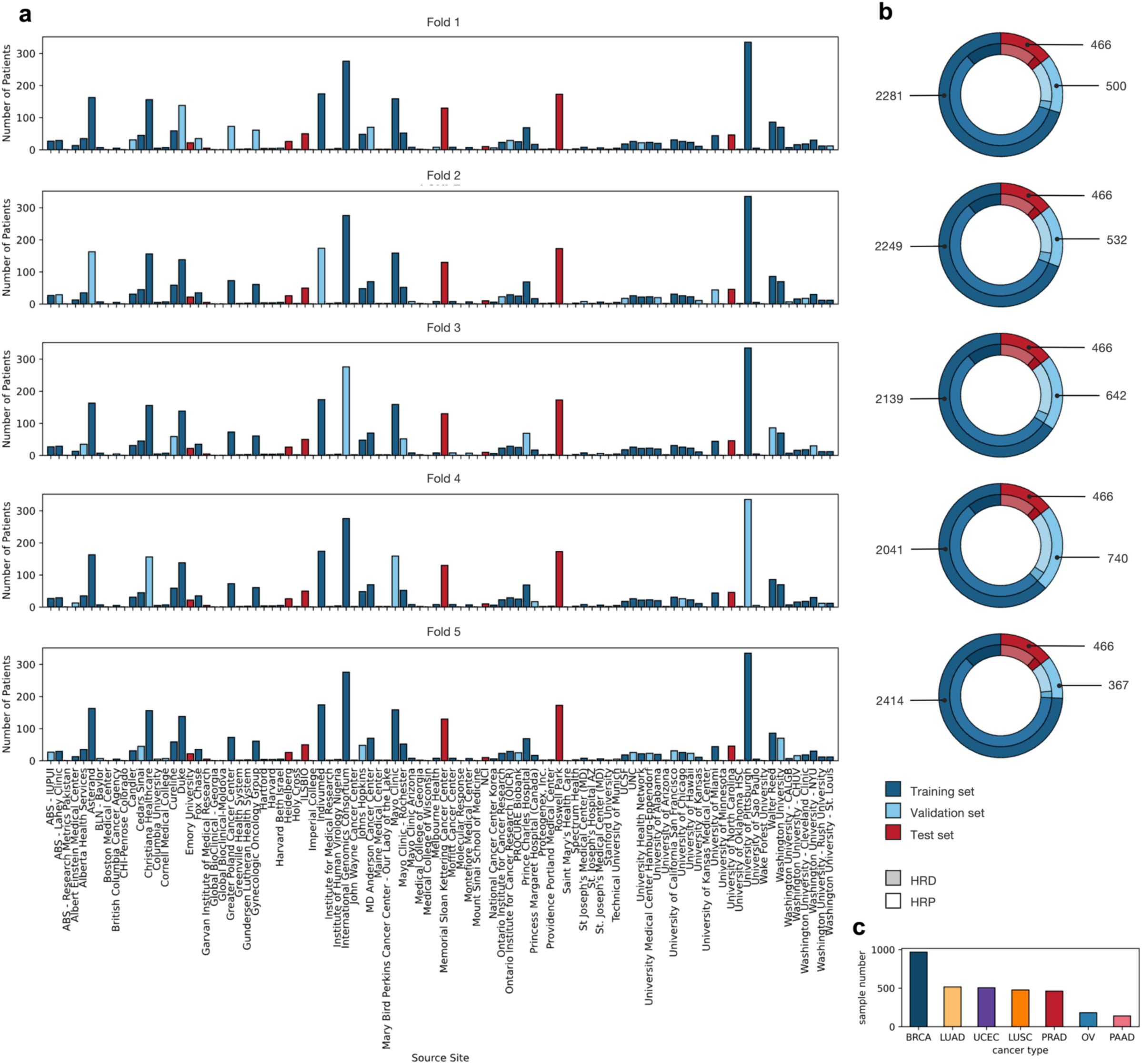
HRD data preparation. **a** Bar chart of site-specific data split for HRD. Dark blue indicates training data, light blue validation data and red the test set. The test set stays the same over all folds. **b** Doughnut plot of the data splits regarding class distribution of HRD vs HRP and number of patients. **c** Bar chart of number of patients per cancer type.

**Supplementary Fig. 3.**
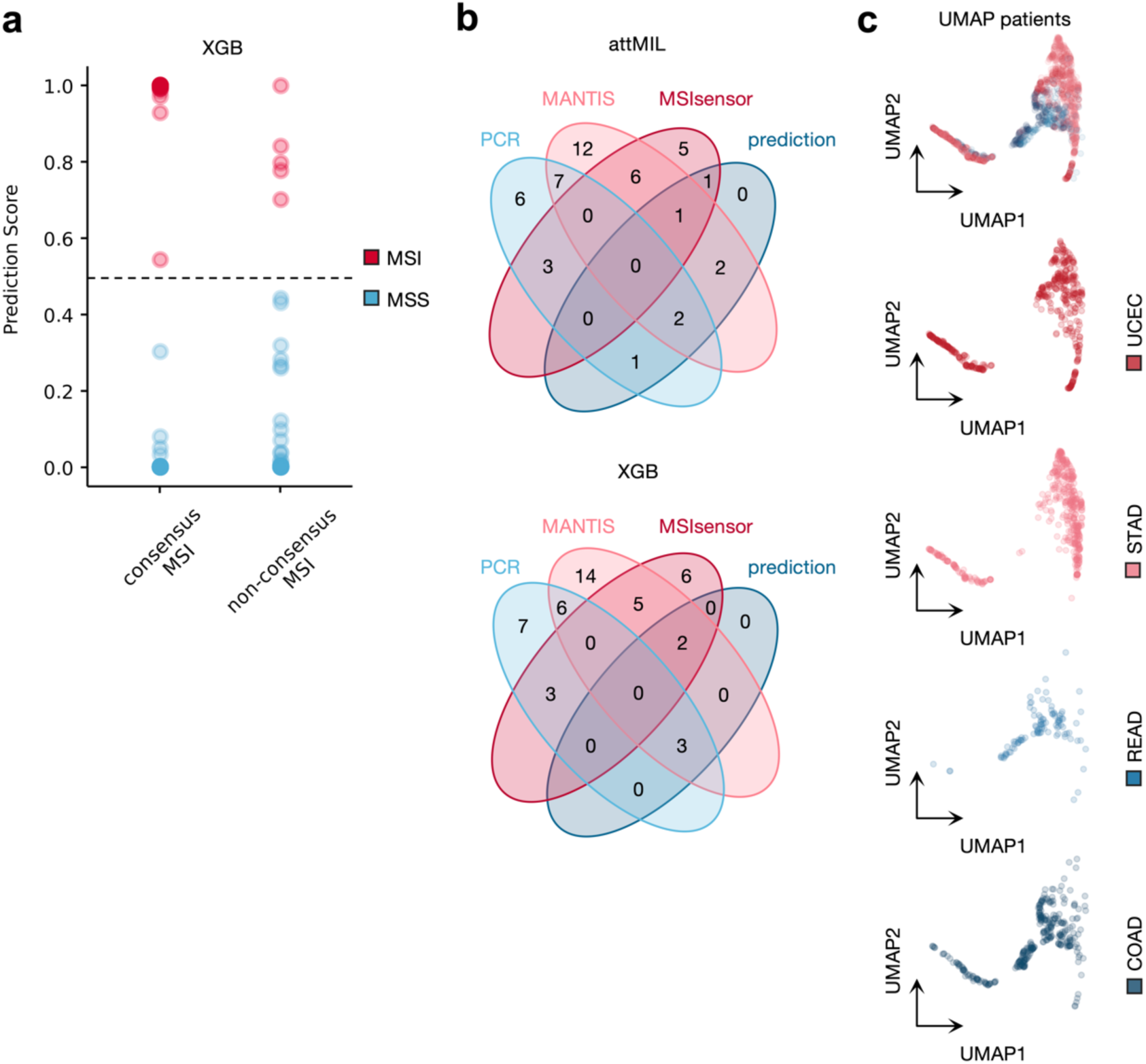
Patient level features of MSI. **a** Prediction scores for consensus and non-consensus patients colored by class (red = MSI, blue = MSS) for XGBoost model. **b** Venn Diagrams of MSI status defined by PCR, MANTIS, MSIsensor and the prediction of the attMIL and XGBoost model. **c** UMAP of patient level features separated by tissue type (dark blue = COAD, blue = READ, rose = STAD, red = UCEC).

**Supplementary Fig. 4.**
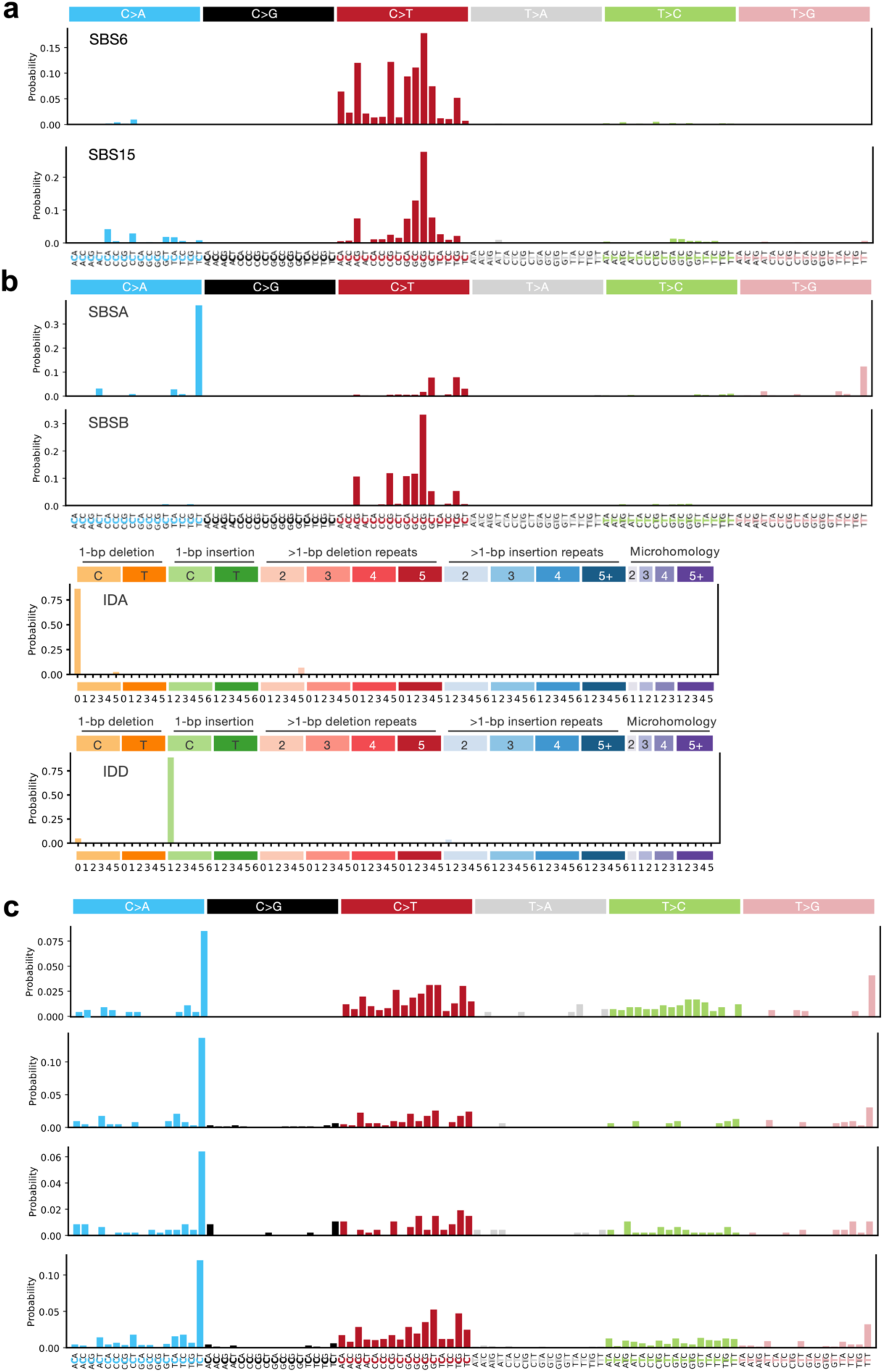
Mutation level features of MSI. **a** COSMIC SBS mutation signatures 6 and 15 associated with MSI. **b** Selected SigProfilerExtractor SBS and Indel mutation signatures similar to mutation catalogues of mutation feature clusters in Fig 2f. **c** SBS mutation catalogues of mutation feature clusters not displayed in Fig. 2f. Only mutation catalogues of the majority mutation class are displayed.

**Supplementary Fig. 5.**
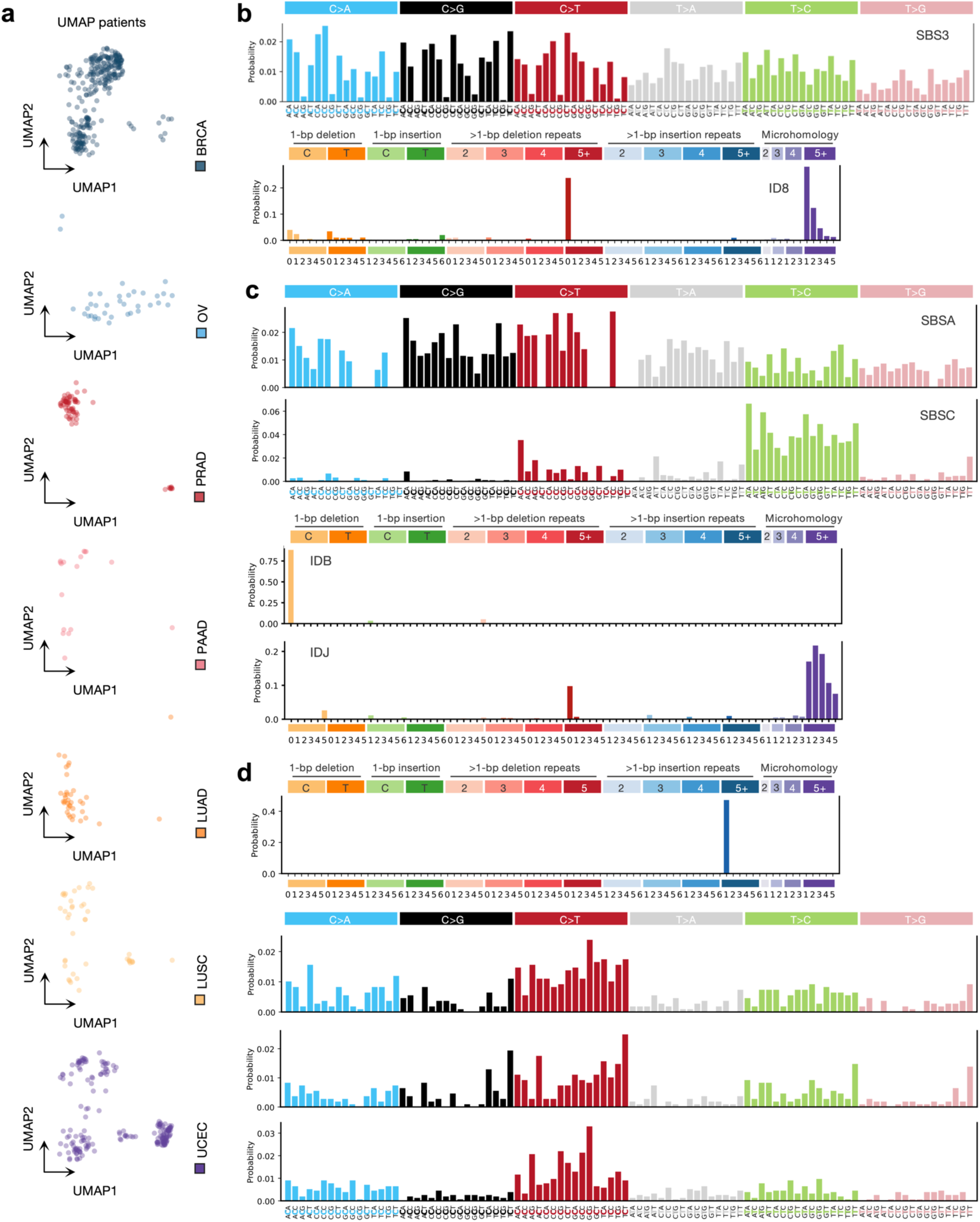
Patient and mutation level features of HRD. **a** UMAP of patient level features separated by tissue type (dark blue = BRCA, blue = OV, rose = PAAD, red = PRAD, orange = LUAD, yellow = LUSC, purple = UCEC). **b** COSMIC SBS mutation signature 3 and indel signature 8 associated with HRD and NHEJ. **c** Selected SigProfilerExtractor SBS and Indel mutation signatures similar to mutation catalogues of mutation feature clusters in Fig. 3c. **d** SBS and indel mutation catalogues of mutation feature clusters not displayed in Fig. 3c. Only mutation catalogues of the majority mutation class are displayed.

**Supplementary Fig. 6.**
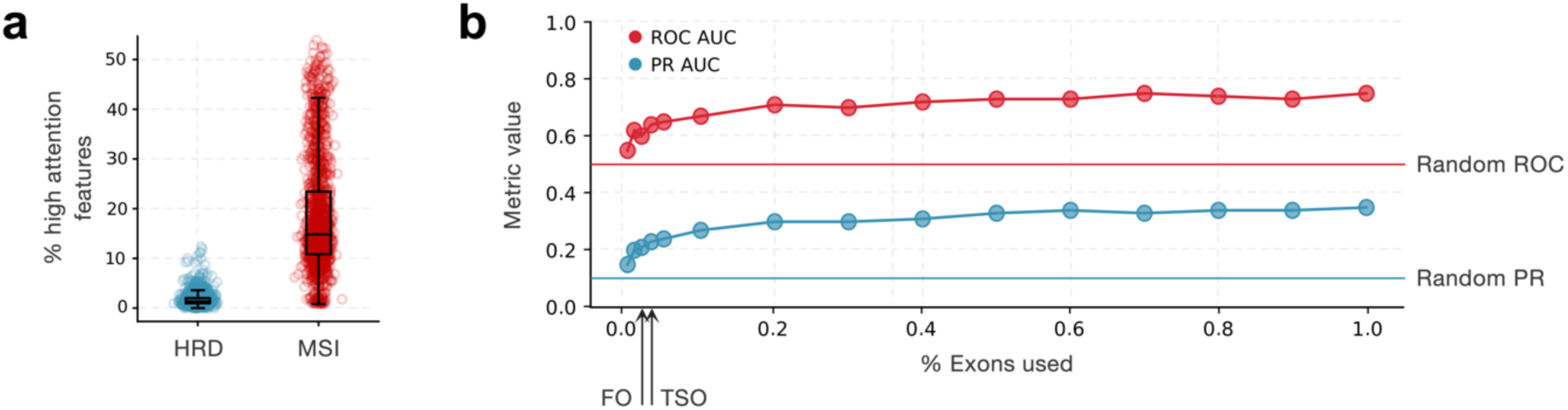
HRD and predictions from somatic mutations only. **a** Comparison of high attention mutations in relation to the total of mutations per patient between MSI and HRD tasks. **b** ROC AUC and PR AUC of the attMIL model on different levels of exome. TruSight Oncology (TSO) and Foundation One (FO) panel reduction indicated by red and blue lines. Random baseline of ROC AUC and PR AUC indicated.

## Supplementary Tables

**Supplementary Table. 1.** MSI patient counts in TCGA. MSI and MSS cases in the TCGA data by cancer type and in total.

|  | COAD | READ | STAD | UCEC | total |
| --- | --- | --- | --- | --- | --- |
| MSI cases | 59 | 3 | 82 | 136 | 280 |
| MSS cases | 279 | 107 | 350 | 314 | 1050 |

**Supplementary Table. 2.**
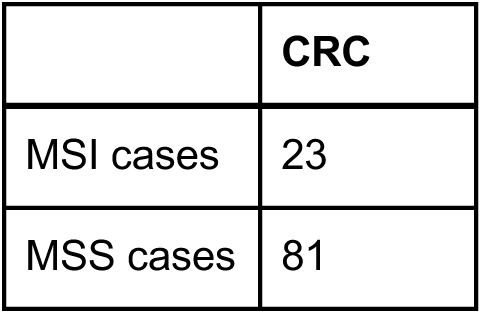
MSI patient counts in CPTAC. MSI and MSS cases in CPTAC CRC.

**Supplementary Table. 3.** MSI patient counts in GENIE. MSI and MSS cases in the GENIE CRC.

|  | CRC |
| --- | --- |
| MSI cases | 25 |
| MSS cases | 296 |

**Supplementary Table. 4.** HRD patient counts in TCGA. HRD and HRP cases in the TCGA data by cancer type and in total.

|  | BRCA | OV | PAAD | PRAD | UCEC | LUAD | LUSC | total |
| --- | --- | --- | --- | --- | --- | --- | --- | --- |
| HRD cases | 180 | 78 | 4 | 5 | 53 | 80 | 164 | 590 |
| HRP cases | 787 | 104 | 135 | 457 | 451 | 436 | 313 | 2657 |

**Supplementary Table. 5.** HRD patient counts in CPTAC. HRD and HRP cases in the CPTAC data by cancer type and in total.

|  | PAAD | LUAD | LUSC | UCE | total |
| --- | --- | --- | --- | --- | --- |
| HRD cases | 2 | 5 | 19 | 1 | 27 |
| HRP cases | 112 | 76 | 70 | 96 | 354 |

**Supplementary Table. 6.** HRD patient counts in GeneseeqPrime® C2. HRD and HRP cases in the CPTAC data by cancer type and in total

|  | OV |
| --- | --- |
| HRD cases | 55 |
| HRP cases | 30 |

**Supplementary Table. 7.**
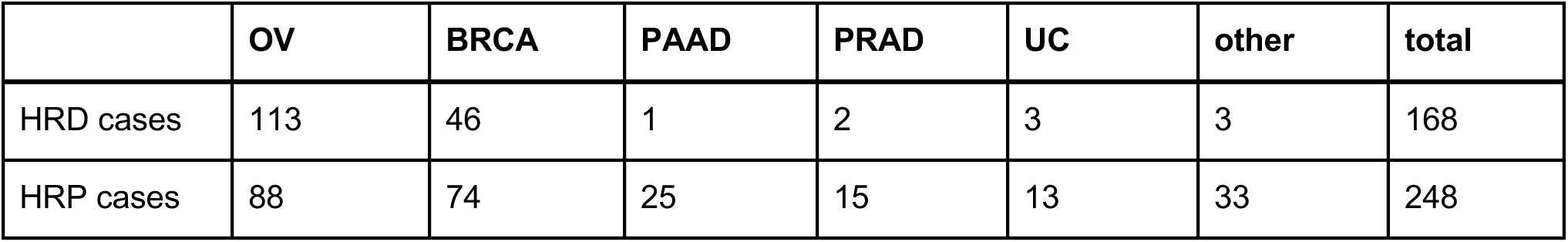
HRD patient counts in GeneseeqPrime® C3. HRD and HRP cases in the CPTAC data by cancer type and in total

|  | OV | BRCA | PAAD | PRAD | UC | other | total |
| --- | --- | --- | --- | --- | --- | --- | --- |
| HRD cases | 113 | 46 | 1 | 2 | 3 | 3 | 168 |
| HRP cases | 88 | 74 | 25 | 15 | 13 | 33 | 248 |

**Supplementary Table. 8.** Comparison of ML Algorithms predicting HRD. Performance values on the full CPTAC external test dataset of the Logistic Regression (LR), Support Vector Machine (SVM), Random Forrest (RF), XGBoost. Best performances are underlined. The p-values were established on the F1 score of the best performing model and all other models.

|  | accuracy | F1 score | ROC AUC | PR AUC | sensitivity | specificity | p-value |
| --- | --- | --- | --- | --- | --- | --- | --- |
| LR | 0.78 ± 0.03 | 0.52 ± 0.06 | 0.83 ± 0.04 | 0.53 ± 0.07 | 0.71 ± 0.07 | <u>0.86 ± 0.05</u> | 0.13 |
| SVM | 0.72 ± 0.01 | 0.4 ± 0.03 | 0.73 ± 0.04 | 0.41 ± 0.04 | 0.68 ± 0.12 | 0.77 ± 0.09 | 0.06 |
| RF | 0.87 ± 0.01 | <u>0.56 ± 0.04</u> | 0.92 ± 0.01 | 0.6 ± 0.03 | 0.93 ± 0.02 | 0.81 ± 0.04 | 1.0 |
| XGBoost | <u>0.88 ± 0.01</u> | <u>0.56 ± 0.03</u> | <u>0.94 ± 0.01</u> | <u>0.69 ± 0.08</u> | <u>0.95 ± 0.03</u> | 0.81 ± 0.03 | - |

**Supplementary Table. 9.** Comparison of CNV encoders predicting HRD. Performance values on the CPTAC external dataset of different encoders: Multi-Layer Perceptron (MPL), Auto Encoder (AU), Variational Auto Encoder (VAE), Masked Auto Encoder (MAE), Masked Language Modelling with Bert (MLM Bert), Masked Language Modelling with Hyena (MLM Hyena). Best performances are underlined. The p-values were established on the F1 score of the best performing model and all other models.

|  | accuracy | F1 score | ROC AUC | PR AUC | sensitivity | specificity | p-value |
| --- | --- | --- | --- | --- | --- | --- | --- |
| MLP | 0.84 ± 0.01 | 0.54 ± 0.03 | 0.88 ± 0.02 | 0.61 ± 0.05 | 0.85 ± 0.04 | 0.82 ± 0.04 | 0.63 |
| AE | 0.85 ± 0.02 | <u>0.56 ± 0.05</u> | 0.91 ± 0.01 | <u>0.67 ± 0.03</u> | 0.87 ± 0.01 | <u>0.84 ± 0.04</u> | - |
| VAE | 0.85 ± 0.01 | 0.53 ± 0.03 | 0.91 ± 0.0 | 0.64 ± 0.01 | 0.88 ± 0.04 | 0.81 ± 0.04 | 0.44 |
| MAE | 0.86 ± 0.01 | <u>0.56 ± 0.05</u> | <u>0.92 ± 0.01</u> | 0.66 ± 0.03 | 0.88 ± 0.03 | 0.83 ± 0.05 | 1.0 |
| MLM Bert | <u>0.87 ± 0.01</u> | 0.52 ± 0.04 | <u>0.92 ± 0.01</u> | 0.59 ± 0.03 | <u>0.93 ± 0.02</u> | 0.8 ± 0.05 | 0.12 |
| MLM Hyena | <u>0.87 ± 0.01</u> | 0.54 ± 0.02 | 0.91 ± 0.01 | 0.55 ± 0.03 | 0.92 ± 0.03 | 0.82 ± 0.02 | 0.44 |
| CNV attMIL | 0.84 ± 0.02 | 0.49 ± 0.03 | <u>0.92 ± 0.02</u> | 0.58 ± 0.09 | 0.92 ± 0.04 | 0.75 ± 0.03 | 0.06 |

**Supplementary Table. 10.** Comparison of modality fusion predicting HRD. Performance values on the CPTAC external dataset of feature concatenation, cross-attention or intermediate-fusion with cross-attention. Best performances are underlined. The p-values were established on the F1 score of the best performing model and all other models.

|  | accuracy | F1 score | ROC AUC | PR AUC | sensitivity | specificity | p-value |
| --- | --- | --- | --- | --- | --- | --- | --- |
| Concat | 0.86 ± 0.01 | <u>0.61 ± 0.03</u> | 0.92 ± 0.01 | 0.71 ± 0.04 | 0.84 ± 0.05 | <u>0.87 ± 0.03</u> | 0.63 |
| Cross-attention | <u>0.87 ± 0.02</u> | <u>0.61 ± 0.06</u> | <u>0.93 ± 0.02</u> | <u>0.74 ± 0.04</u> | <u>0.89 ± 0.05</u> | 0.85 ± 0.05 | - |
| Intermediate- fusion | 0.85 ± 0.02 | 0.57 ± 0.04 | 0.91 ± 0.02 | 0.72 ± 0.03 | 0.87 ± 0.05 | 0.83 ± 0.05 | 0.06 |

**Supplementary Table. 11.**
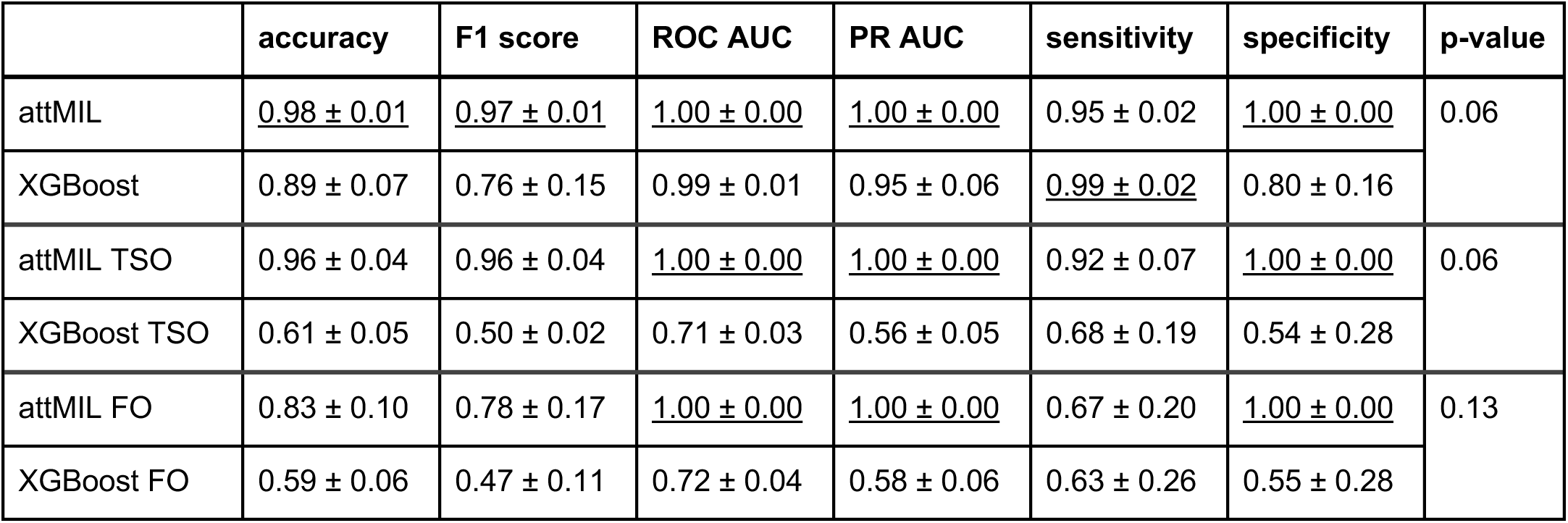
MSI performance metrics in CPTAC. Performance values on the CPTAC external test dataset of attMIL and XGBoost predicting MSI. Best performances are underlined. P-values were generated between pairs of DL/ML models on the F1 score for the same data.

|  | accuracy | F1 score | ROC AUC | PR AUC | sensitivity | specificity | p-value |
| --- | --- | --- | --- | --- | --- | --- | --- |
| attMIL | <u>0.98 ± 0.01</u> | <u>0.97 ± 0.01</u> | <u>1.00 ± 0.00</u> | <u>1.00 ± 0.00</u> | 0.95 ± 0.02 | <u>1.00 ± 0.00</u> | 0.06 |
| XGBoost | 0.89 ± 0.07 | 0.76 ± 0.15 | 0.99 ± 0.01 | 0.95 ± 0.06 | <u>0.99 ± 0.02</u> | 0.80 ± 0.16 |  |
| attMIL TSO | 0.96 ± 0.04 | 0.96 ± 0.04 | <u>1.00 ± 0.00</u> | <u>1.00 ± 0.00</u> | 0.92 ± 0.07 | <u>1.00 ± 0.00</u> | 0.06 |
| XGBoost TSO | 0.61 ± 0.05 | 0.50 ± 0.02 | 0.71 ± 0.03 | 0.56 ± 0.05 | 0.68 ± 0.19 | 0.54 ± 0.28 |  |
| attMIL FO | 0.83 ± 0.10 | 0.78 ± 0.17 | <u>1.00 ± 0.00</u> | <u>1.00 ± 0.00</u> | 0.67 ± 0.20 | <u>1.00 ± 0.00</u> | 0.13 |
| XGBoost FO | 0.59 ± 0.06 | 0.47 ± 0.11 | 0.72 ± 0.04 | 0.58 ± 0.06 | 0.63 ± 0.26 | 0.55 ± 0.28 |  |

**Supplementary Table. 12.** MSI performance metrics in GENIE. Performance values on the GENIE external panel dataset of attMIL and XGBoost predicting MSI. Best performances are underlined. P-value was generated between DL and ML model on the F1 score for the same data.

|  | accuracy | F1 score | ROC AUC | PR AUC | sensitivity | specificity | p-value |
| --- | --- | --- | --- | --- | --- | --- | --- |
| attMIL | <u>0.68 ± 0.06</u> | <u>0.52 ± 0.13</u> | <u>0.85 ± 0.01</u> | <u>0.68 ± 0.03</u> | 0.37 ± 0.12 | <u>1.00 ± 0.00</u> | 0.06 |
| XGBoost | 0.58 ± 0.05 | 0.21 ± 0.05 | 0.64 ± 0.05 | 0.25 ± 0.07 | <u>0.52 ± 0.30</u> | 0.63 ± 0.34 |  |
| attMIL (>5) | 0.71 ± 0.07 | 0.57 ± 0.13 | 0.88 ± 0.01 | 0.76 ± 0.03 | 0.42 ± 0.14 | 1.00 ± 0.00 | - |
| attMIL (>10) | 0.73 ± 0.07 | 0.60 ± 0.14 | 0.92 ± 0.02 | 0.84 ± 0.03 | 0.46 ± 0.15 | 1.00 ± 0.00 | - |

**Supplementary Table. 13.**
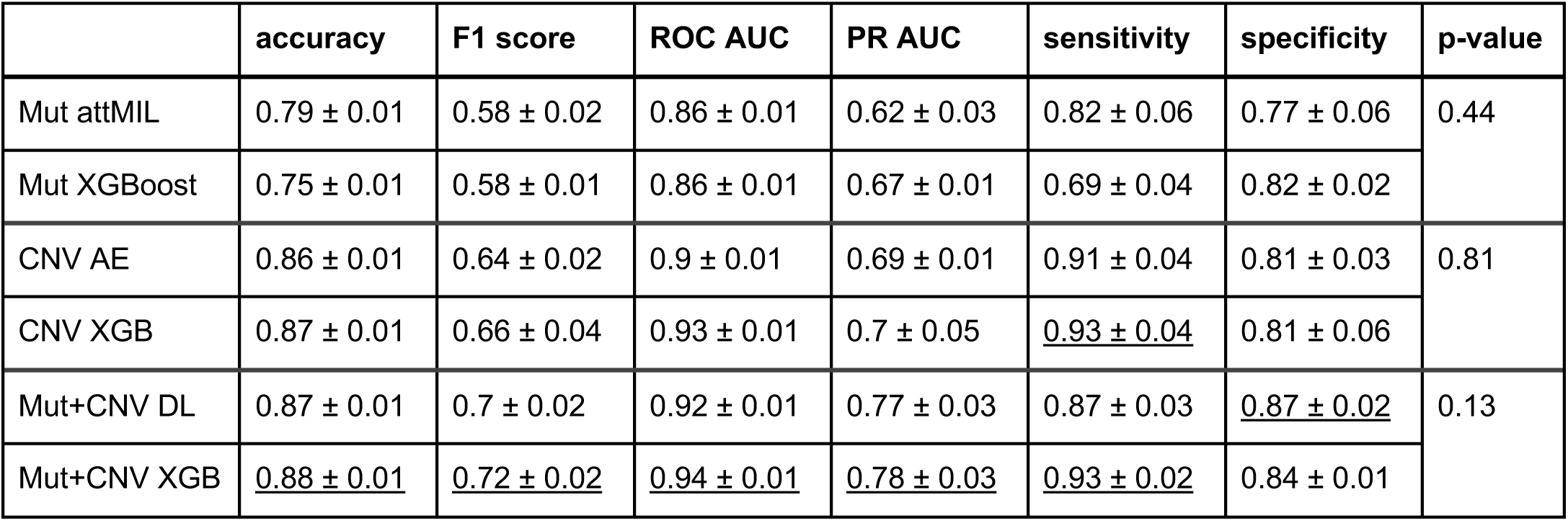
HRD performance metrics in TCGA. Performance values on the TCGA test dataset predicting HRD of single and multi-modality models. Best performances are underlined. P-values were generated between pairs of DL/ML models on the F1 score for the same data.

**Supplementary Table. 14.**
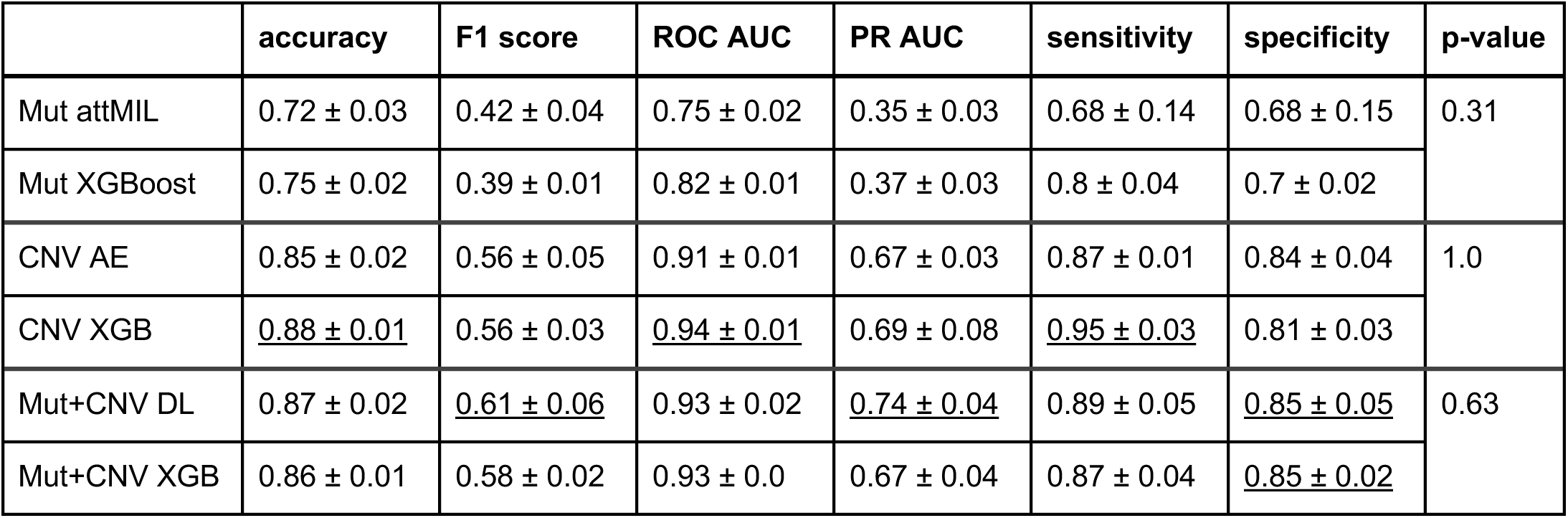
HRD performance metrics in CPTAC. Performance values on the full CPTAC external test dataset predicting HRD of single and multi-modality models. P-values were generated between pairs of DL/ML models on the F1 score for the same data.

|  | accuracy | F1 score | ROC AUC | PR AUC | sensitivity | specificity | p-value |
| --- | --- | --- | --- | --- | --- | --- | --- |
| Mut attMIL | 0.72 ± 0.03 | 0.42 ± 0.04 | 0.75 ± 0.02 | 0.35 ± 0.03 | 0.68 ± 0.14 | 0.68 ± 0.15 | 0.31 |
| Mut XGBoost | 0.75 ± 0.02 | 0.39 ± 0.01 | 0.82 ± 0.01 | 0.37 ± 0.03 | 0.8 ± 0.04 | 0.7 ± 0.02 |  |
| CNV AE | 0.85 ± 0.02 | 0.56 ± 0.05 | 0.91 ± 0.01 | 0.67 ± 0.03 | 0.87 ± 0.01 | 0.84 ± 0.04 | 1.0 |
| CNV XGB | <u>0.88 ± 0.01</u> | 0.56 ± 0.03 | <u>0.94 ± 0.01</u> | 0.69 ± 0.08 | <u>0.95 ± 0.03</u> | 0.81 ± 0.03 |  |
| Mut+CNV DL | 0.87 ± 0.02 | <u>0.61 ± 0.06</u> | 0.93 ± 0.02 | <u>0.74 ± 0.04</u> | 0.89 ± 0.05 | <u>0.85 ± 0.05</u> | 0.63 |
| Mut+CNV XGB | 0.86 ± 0.01 | 0.58 ± 0.02 | 0.93 ± 0.0 | 0.67 ± 0.04 | 0.87 ± 0.04 | <u>0.85 ± 0.02</u> |  |

**Supplementary Table. 15.** HRD performance metrics in GeneseeqPrime®. Performance values on the full CPTAC external test dataset predicting HRD of single and multi-modality models. P-values were generated between pairs of DL/ML models on the F1 score for the same data.

|  | accuracy | F1 score | ROC AUC | PR AUC | sensitivity | specificity | p-value |
| --- | --- | --- | --- | --- | --- | --- | --- |
| XGBoost trained on TCGA | 0.58 ± 0.06 | 0.42 ± 0.27 | 0.72 ± 0.09 | 0.83 ± 0.06 | 0.35 ± 0.29 | 0.81 ± 0.21 | 0.19 |
| AE trained on TCGA | 0.75 ± 0.09 | 0.76 ± 0.16 | 0.89 ± 0.03 | 0.93 ± 0.02 | 0.72 ± 0.23 | 0.78 ± 0.22 |  |
| XGBoost calibrated | 0.66 ± 0.06 | 0.62 ± 0.10 | 0.75 ± 0.04 | 0.84 ± 0.04 | 0.51 ± 0.11 | 0.81 ± 0.04 | 0.13 |
| AE calibrated | 0.78 ± 0.04 | 0.79 ± 0.05 | 0.89 ± 0.02 | 0.93 ± 0.01 | 0.71 ± 0.07 | 0.85 ± 0.05 |  |
| XGBoost trained on GeneseeqPrime® | 0.86 ± 0.02 | 0.90 ± 0.02 | 0.96 ± 0.02 | 0.98 ± 0.01 | 0.91 ± 0.06 | 0.81 ± 0.06 | 0.44 |
| AE trained on GeneseeqPrime® | 0.90 ± 0.01 | 0.93 ± 0.01 | 0.96 ± 0.01 | 0.98 ± 0.01 | 0.93 ± 0.02 | 0.88 ± 0.04 |  |

## Supplementary Methods

### Feature Catalogues for ML model

To generate the SBS mutation catalogues, patient mutations were grouped by 96 SBS features, encompassing six mutation classes: C>A, C>G, C>T, T>A, T>C, T>G (including the reverse complement). Each class was further stratified by the adjacent 3’ and 5’ nucleotides. Since there are four possible options for the 3’ and 5’ positions, this results in 6 * 4 * 4 = 96 features. SBS mutations are displayed as, for example, a C>T mutation with an upstream G and a downstream C (GC>TC). Indel mutations can be read as first the indel pattern length and then the number of repetitions. For example, an insertion pattern of at least five nucleotides without a repetition (5InsRep0) or a mononucleotide deletion of a C with the length one (1DelC1).

For indel feature generation, mutations were grouped according to insertion or deletion length, repetition pattern, and a possible homology to flanking sites. For mononucleotide deletions of C or T (including complementary base), two classes were created, along with two classes for corresponding mononucleotide insertions. Additionally, four classes were defined based on repetition patterns for deletions and insertions. Each of these 12 initial subclasses was then subdivided into six subclasses based on the number of repeats deleted or inserted, resulting in 72 subclasses. Finally, four microhomology classes were assigned, describing mutations in which either the 3’ or 5’ flanking site has partial homology to the deletion sequence. These classes were further subdivided based on homology length, adding 11 additional classes to the 72 remaining classes, resulting in a total of 83 indel classes. Microhomology mutations can be read as first the deletion length and then the length of the microhomology. For example, a deletion of at least five nucleotides containing a microhomology of two nucleotides to at least one of the flanking sites (5+DelMH2).

### Mutation Signatures

Mutation signatures provide an alternative feature representation to mutation catalogues by summarizing mutational processes, including signatures associated with dMMR and HRD, via per-sample exposure profiles. We therefore used SigProfilerMatrixGenerator [73] and SigProfilerAssignment [74] to extract 99 SBS and 23 indel signature exposures, normalized exposures per patient, and used them as input features for an XGBoost model (**Supplementary Results**).

### Multiple Instance Learning

Multiple Instance Learning (MIL) is a deep learning framework designed to handle scenarios where labels are associated with sets of instances, referred to as “bags,” rather than individual instances. In MIL, only a subset of instances within a bag contribute to the overall class prediction, while others may be irrelevant or unrelated. This makes MIL particularly suitable for contexts where identifying which instances are informative is inherently challenging or unknown [52]

Here, we applied MIL to model patient-level predictions, where each patient is represented as a “bag” containing multiple somatic mutations (“instances”) [32,39]. Not all mutations contribute to the phenotype of interest (e.g., MSI or HRD), as some mutations may not be associated with the underlying mutational mechanisms driving these biomarkers. MIL allows the model to autonomously learn which mutations are most relevant by assigning attention weights to individual instances, rather than relying on predefined rules or human-curated knowledge.

### Masked-language modelling

Masked language modelling (MLM) was used to train CNV encoders in a self-supervised manner. Each patient was represented by a fixed-length vector of gene-level CNV values (19,444 genes, log₂ copy-number ratios). Continuous CNV values were discretised into 62 bins, and each bin was assigned a token ID. Thus, each gene corresponds to one discrete token, and the CNV profile of a patient forms a sequence of 19,444 tokens. Additional special tokens (e.g. [CLS], [MASK], [PAD]) were included as in standard MLM setups. [18,75]

During training, 15% of tokens were randomly selected and replaced by a [MASK] token. The model received the masked sequence as input and was trained to predict the original token at the masked positions. This objective encourages the model to learn dependencies between copy-number states at different genomic positions and to form internal representations of typical CNV patterns across the genome. Training used a token-level cross-entropy loss computed only at masked positions, averaged across masked tokens in the batch.

We implemented this task using a BERT-style transformer encoder [53]. Each token ID is mapped to a continuous vector (“embedding”), and several self-attention layers update these embeddings by allowing each token to attend to all other tokens in the sequence in a bidirectional manner. This enables the model to use both upstream and downstream CNV context when reconstructing masked genes. However, self-attention has quadratic complexity in sequence length, which makes standard transformers computationally expensive for long CNV sequences (19,944 positions). In practice, this limited the feasible embedding dimension in BERT-based models on our hardware.

To mitigate these constraints, we additionally trained a Hyena-based encoder [54], which replaces full self-attention with structured convolutional operators that approximate long-range dependencies with lower computational and memory cost. This allowed us to handle the same CNV sequence length with larger embedding sizes under the same resource budget, while using the same MLM objective (masking and reconstructing binned CNV tokens).

### Pseudo-panel MSI mutation inputs and bag-size sensitivity analyses

For MSI analyses, pseudo-panel datasets were generated by restricting exome-derived somatic small variants (SNVs/indels) to the gene sets of common targeted panels (TSO and FoundationOne). Variants outside the respective gene list were removed. These pseudo-panels were used to study MSI prediction under reduced sequencing coverage while keeping upstream processing consistent with the exome cohorts.

Because pseudo-panel restriction substantially reduces the number of mutation instances per sample, we assessed the sensitivity of the attention-based MIL model to changes in bag size. In addition to the upsampling procedure described in the main Methods, we implemented a fixed-bag variant in which each sample is represented by exactly *κ* mutation instances per forward pass. This analysis was designed to evaluate whether enforcing a constant bag size yields comparable performance without relying on instance repetition. For samples with at least *κ* instances, *κ* instances were sampled uniformly without replacement. For samples with fewer than *κ* instances, instances were sampled with replacement until *κ* were obtained.

Fixed-bag models were trained and evaluated with the same pseudo-panel inputs to assess whether enforcing a constant instance count improves robustness under panel-like sparsity. We used bag sizes *κ* = 20 and *κ* = 80, since the mean mutation count of GENIE and CPTAC pseudo-panel is 20 and 80 for TCGA.

To separate training-distribution effects from evaluation constraints, we trained MIL models either on exome-wide mutation inputs and evaluated them on pseudo-panels, or trained directly on pseudo-panel inputs. Performance was assessed on internal test splits and on independent external cohorts.

## Supplementary Results

### Mutation Signatures vs Mutational Catalogues

In our comparisons, signature-based features did not improve MSI prediction relative to catalogue features and performed worse for HR. We therefore retained the catalogue representation as the primary ML input throughout the manuscript. In addition, because part of our evaluation focuses on panel and panel-like settings, we did not use signatures as a default feature set, as signature extraction becomes less stable when mutation counts are low.

#### Comparison of mutation catalogues vs mutation signatures for MSI

Performance values on the full CPTAC external test dataset of XGBoost applied on SBS and indel mutational signatures (112 features) and mutational catalogues (179 features). Best performances are underlined. The p-value was generated on the F1 score for the same data.

|  | accuracy | F1 score | ROC AUC | PR AUC | sensitivity | specificity | p-value |
| --- | --- | --- | --- | --- | --- | --- | --- |
| Mutation signatures | $0.87 \pm 0.08$ | <u>$0.82 \pm 0.11$</u> | $0.98 \pm 0.02$ | <u>$0.95 \pm 0.02$</u> | $0.77 \pm 0.15$ | <u>$0.98 \pm 0.03$</u> | 0.44 |
| Mutation catalogues | <u>$0.89 \pm 0.07$</u> | $0.76 \pm 0.15$ | <u>$0.99 \pm 0.01$</u> | <u>$0.95 \pm 0.06$</u> | <u>$0.99 \pm 0.02$</u> | $0.80 \pm 0.16$ | |

#### Comparison of mutation catalogues vs mutation signatures for HRD

Performance values on the full CPTAC external test dataset of XGBoost applied on SBS and indel mutational signatures and mutational catalogues. Best performances are underlined. The p-value was generated on the F1 score for the same data.

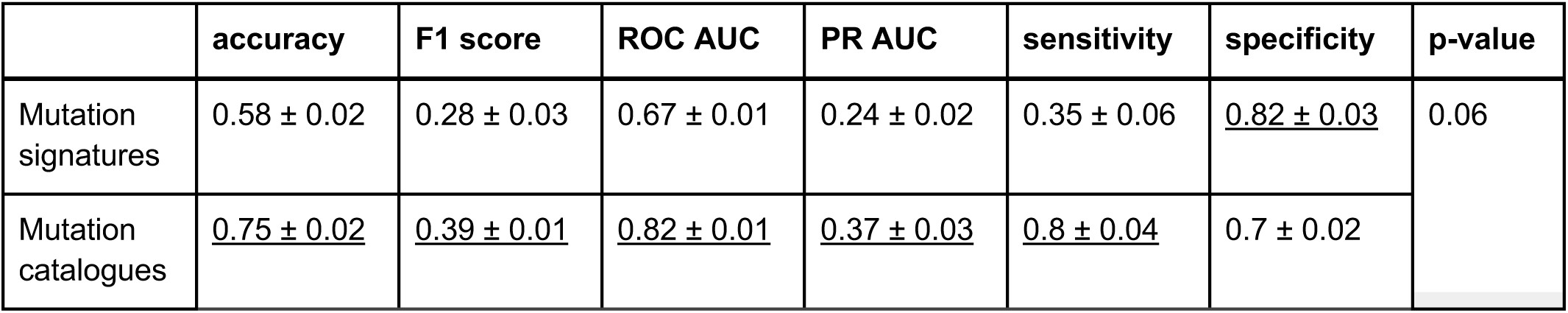

|  | accuracy | F1 score | ROC AUC | PR AUC | sensitivity | specificity | p-value |
| --- | --- | --- | --- | --- | --- | --- | --- |
| Mutation signatures | $0.58 \pm 0.02$ | $0.28 \pm 0.03$ | $0.67 \pm 0.01$ | $0.24 \pm 0.02$ | $0.35 \pm 0.06$ | <u>$0.82 \pm 0.03$</u> | 0.06 |
| Mutation catalogues | <u>$0.75 \pm 0.02$</u> | <u>$0.39 \pm 0.01$</u> | <u>$0.82 \pm 0.01$</u> | <u>$0.37 \pm 0.03$</u> | <u>$0.8 \pm 0.04$</u> | $0.7 \pm 0.02$ | |

### Sensitivity of MSI MIL models to pseudo-panel sparsity and bag size

As shown in the main Results, MIL achieved near-ceiling performance for MSI on exome-wide inputs and retained strong performance under pseudo-panel restriction with the upsampling procedure described in Methods, while performance decreased on GENIE. To further probe whether this behavior reflects training distribution or bag-size effects, we performed two additional analyses.

First, we trained MIL directly on TCGA pseudo-panels and evaluated it on external pseudo-panel cohorts. Internal performance remained high (TCGA: F1 0.97 ± 0.02, PR-AUC 1.00 ± 0.00), while external performance decreased relative to TCGA, with strong transfer to CPTAC (F1 0.85 ± 0.01, PR-AUC 0.98 ± 0.02) and lower performance on GENIE (PR-AUC 0.66 ± 0.07). Across cohorts, PR-AUC was more stable than threshold-dependent F1, consistent with threshold sensitivity under cohort and assay differences.

Second, we evaluated a fixed-bag MIL variant to enforce a constant instance count per sample. In this configuration (*κ*=80), performance decreased already on TCGA and deteriorated further on external cohorts (TCGA: F1 0.82 ± 0.04, PR-AUC 0.85 ± 0.03; CPTAC: F1 0.30 ± 0.19, PR-AUC 0.60 ± 0.11; GENIE: F1 0.12 ± 0.09, PR-AUC 0.16 ± 0.05), indicating that this fixed-bag setting was not suitable for sparse pseudo-panel mutation inputs in our implementation.

Overall, these analyses indicate that MSI prediction from panel-like mutation inputs is sensitive to mutation burden and cohort differences, and that bag-size control strategies can substantially influence thresholded performance.

#### Performance of MSI MIL models on TCGA under exome-wide and pseudo-panel settings

Performance values for four configurations: (i) exome-wide training/evaluation, (ii) exome-trained model evaluated on (pseudo-)panels (with bag-size upsampling as in Methods), (iii) pseudo-panel-trained model on (pseudo-)panels (with bag-size upsampling as in Methods), and (iv) pseudo-panel-trained fixed-bag MIL model (constant bag size *κ*). Best performances are underlined. P-values were generated between the best performing model and all others on the F1 score for the same data.

|  | accuracy | F1 score | ROC AUC | PR AUC | sensitivity | specificity | p-value |
| --- | --- | --- | --- | --- | --- | --- | --- |
| i | <u>$1.00 \pm 0.00$</u> | <u>$1.00 \pm 0.00$</u> | <u>$1.00 \pm 0.00$</u> | <u>$1.00 \pm 0.00$</u> | <u>$1.00 \pm 0.00$</u> | <u>$1.00 \pm 0.00$</u> | - |
| ii | $0.93 \pm 0.09$ | $0.90 \pm 0.10$ | <u>$1.00 \pm 0.01$</u> | $0.99 \pm 0.01$ | $0.86 \pm 0.19$ | $0.99 \pm 0.02$ | 0.07 |
| iii | $0.99 \pm 0.01$ | $0.97 \pm 0.02$ | $1.00 \pm 0.00$ | $1.00 \pm 0.00$ | $1.00 \pm 0.00$ | $0.99 \pm 0.01$ | 0.07 |
| iv $\kappa = 20$ | $0.93 \pm 0.01$ | $0.81 \pm 0.04$ | $0.96 \pm 0.02$ | $0.84 \pm 0.11$ | $0.98 \pm 0.02$ | $0.88 \pm 0.04$ | 0.06 |
| iv $\kappa = 80$ | $0.92 \pm 0.02$ | $0.82 \pm 0.04$ | $0.96 \pm 0.06$ | $0.85 \pm 0.03$ | $0.92 \pm 0.06$ | $0.91 \pm 0.04$ | 0.06 |

#### Performance of MSI MIL models on CPTAC under exome-wide and pseudo-panel settings

Performance values for four configurations: (i) exome-wide training/evaluation, (ii) exome-trained model evaluated on pseudo-panels (with bag-size upsampling as in Methods), (iii) pseudo-panel-trained model, and (iv) pseudo-panel-trained fixed-bag MIL model (constant bag size). Best performances are underlined. P-values were generated between the best performing model and all others on the F1 score for the same data.

|  | accuracy | F1 score | ROC AUC | PR AUC | sensitivity | specificity | p-value |
| --- | --- | --- | --- | --- | --- | --- | --- |
| i | <u>0.98 ± 0.01</u> | <u>0.97 ± 0.01</u> | <u>1.00 ± 0.00</u> | <u>1.00 ± 0.00</u> | <u>0.95 ± 0.02</u> | <u>1.00 ± 0.00</u> | - |
| ii | 0.83 ± 0.10 | 0.78 ± 0.17 | <u>1.00 ± 0.00</u> | <u>1.00 ± 0.00</u> | 0.67 ± 0.20 | <u>1.00 ± 0.00</u> | 0.06 |
| iii | 0.88 ± 0.06 | 0.85 ± 0.01 | 0.99 ± 0.01 | 0.98 ± 0.02 | 0.76 ± 0.11 | <u>1.00 ± 0.01</u> | 0.06 |
| iv $\kappa = 20$ | 0.66 ± 0.06 | 0.46 ± 0.14 | 0.91 ± 0.03 | 0.73 ± 0.12 | 0.35 ± 0.13 | 0.97 ± 0.03 | 0.06 |
| iv $\kappa = 80$ | 0.59 ± 0.07 | 0.30 ± 0.19 | 0.86 ± 0.03 | 0.60 ± 0.11 | 0.22 ± 0.17 | 0.96 ± 0.04 | 0.06 |

#### Performance of MSI MIL models on GENIE

Performance values for four configurations: (ii) exome-trained model evaluated on panel data (with bag-size upsampling as in Methods), (iii) pseudo-panel-trained model evaluated on panel data, and (iv) pseudo-panel-trained fixed-bag MIL model (constant bag size) evaluated on panel data. Best performances are underlined. P-values were generated between the best performing model and all others on the F1 score for the same data.

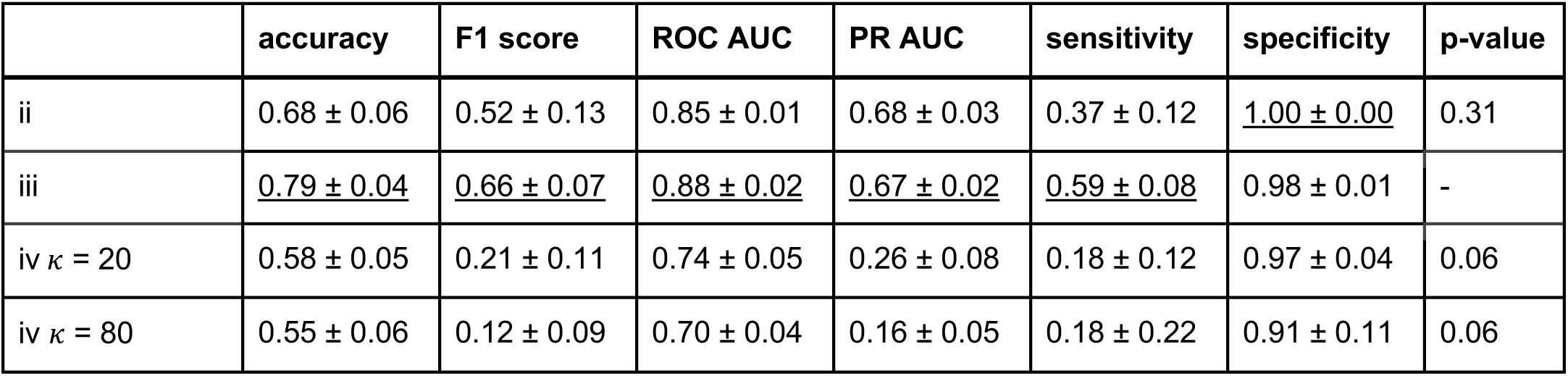

|  | accuracy | F1 score | ROC AUC | PR AUC | sensitivity | specificity | p-value |
| --- | --- | --- | --- | --- | --- | --- | --- |
| ii | 0.68 ± 0.06 | 0.52 ± 0.13 | 0.85 ± 0.01 | 0.68 ± 0.03 | 0.37 ± 0.12 | <u>1.00 ± 0.00</u> | 0.31 |
| iii | <u>0.79 ± 0.04</u> | <u>0.66 ± 0.07</u> | <u>0.88 ± 0.02</u> | <u>0.67 ± 0.02</u> | <u>0.59 ± 0.08</u> | 0.98 ± 0.01 | - |
| iv $\kappa = 20$ | 0.58 ± 0.05 | 0.21 ± 0.11 | 0.74 ± 0.05 | 0.26 ± 0.08 | 0.18 ± 0.12 | 0.97 ± 0.04 | 0.06 |
| iv $\kappa = 80$ | 0.55 ± 0.06 | 0.12 ± 0.09 | 0.70 ± 0.04 | 0.16 ± 0.05 | 0.18 ± 0.22 | 0.91 ± 0.11 | 0.06 |

## References

1. Andre F, Filleron T, Kamal M, Mosele F, Arnedos M, Dalenc F, et al. Genomics to select treatment for patients with metastatic breast cancer. Nature. 2022;610:343–8.

2. Mateo J, Steuten L, Aftimos P, André F, Davies M, Garralda E, et al. Delivering precision oncology to patients with cancer. Nat Med. 2022;28:658–65.

3. Hodder A, Leiter SM, Kennedy J, Addy D, Ahmed M, Ajithkumar T, et al. Benefits for children with suspected cancer from routine whole-genome sequencing. Nat Med. 2024;30:1905–12.

4. Pleasance ED, Cheetham RK, Stephens PJ, McBride DJ, Humphray SJ, Greenman CD, et al. A comprehensive catalogue of somatic mutations from a human cancer genome. Nature. 2010;463:191–6.

5. Horak P, Heining C, Kreutzfeldt S, Hutter B, Mock A, Hüllein J, et al. Comprehensive genomic and transcriptomic analysis for guiding therapeutic decisions in patients with rare cancers. Cancer Discov. 2021;11:2780–95.

6. Alexandrov LB, Nik-Zainal S, Wedge DC, Aparicio SAJR, Behjati S, Biankin AV, et al. Signatures of mutational processes in human cancer. Nature. 2013;500:415–21.

7. Gonzalez-Perez A, Perez-Llamas C, Deu-Pons J, Tamborero D, Schroeder MP, Jene-Sanz A, et al. IntOGen-mutations identifies cancer drivers across tumor types. Nat Methods. 2013;10:1081–2.

8. Macintyre G, Goranova TE, De Silva D, Ennis D, Piskorz AM, Eldridge M, et al. Copy number signatures and mutational processes in ovarian carcinoma. Nat Genet. 2018;50:1262–70.

9. Alexandrov LB, Kim J, Haradhvala NJ, Huang MN, Tian Ng AW, Wu Y, et al. The repertoire of mutational signatures in human cancer. Nature. 2020;578:94–101.

10. Li Y, Roberts ND, Wala JA, Shapira O, Schumacher SE, Kumar K, et al. Patterns of somatic structural variation in human cancer genomes. Nature. 2020;578:112–21.

11. Steele CD, Abbasi A, Islam SMA, Bowes AL, Khandekar A, Haase K, et al. Signatures of copy number alterations in human cancer. Nature. 2022;606:984–91.

12. Unger M, Kather JN. Deep learning in cancer genomics and histopathology. Genome Med. 2024;16:44.

13. Unger M, Kather JN. A systematic analysis of deep learning in genomics and histopathology for precision oncology. BMC Med Genomics. 2024;17:48.

14. Kris A. Wetterstrand MS. DNA sequencing costs: Data [Internet]. Genome.gov. NHGRI; 2019 [cited 2024 Sep 9]. Available from: https://www.genome.gov/about-genomics/fact-sheets/DNA-Sequencing-Costs-Data

15. Chiu Y-C, Chen H-IH, Zhang T, Zhang S, Gorthi A, Wang L-J, et al. Predicting drug response of tumors from integrated genomic profiles by deep neural networks. BMC Med Genomics. 2019;12:18.

16. Elmarakeby HA, Hwang J, Arafeh R, Crowdis J, Gang S, Liu D, et al. Biologically informed deep neural network for prostate cancer discovery. Nature. 2021;598:348–52.

17. Ahlmann-Eltze C, Huber W, Anders S. Deep-learning-based gene perturbation effect prediction does not yet outperform simple linear baselines. Nat Methods. 2025;22:1657–61.

18. Gélard M, Richard G, Pierrot T, Cournède P-H. BulkRNABert: Cancer prognosis from bulk RNA-seq based language models [Internet]. bioRxiv. 2024. Available from: 10.1101/2024.06.18.599483

19. Benkirane H, Pradat Y, Michiels S, Cournède P-H. CustOmics: A versatile deep-learning based strategy for multi-omics integration. PLoS Comput Biol. 2023;19:e1010921.

20. Wang F-A, Zhuang Z, Gao F, He R, Zhang S, Wang L, et al. TMO-Net: an explainable pretrained multi-omics model for multi-task learning in oncology. Genome Biol. 2024;25:149.

21. Le DT, Durham JN, Smith KN, Wang H, Bartlett BR, Aulakh LK, et al. Mismatch repair deficiency predicts response of solid tumors to PD-1 blockade. Science. 2017;357:409–13.

22. Germano G, Lamba S, Rospo G, Barault L, Magrì A, Maione F, et al. Inactivation of DNA repair triggers neoantigen generation and impairs tumour growth. Nature. 2017;552:116–20.

23. Chopra N, Tovey H, Pearson A, Cutts R, Toms C, Proszek P, et al. Homologous recombination DNA repair deficiency and PARP inhibition activity in primary triple negative breast cancer. Nat Commun. 2020;11:2662.

24. Polak P, Kim J, Braunstein LZ, Karlic R, Haradhavala NJ, Tiao G, et al. A mutational signature reveals alterations underlying deficient homologous recombination repair in breast cancer. Nat Genet. 2017;49:1476–86.

25. Moore Kathleen, Colombo Nicoletta, Scambia Giovanni, Kim Byoung-Gie, Oaknin Ana, Friedlander Michael, et al. Maintenance Olaparib in Patients with Newly Diagnosed Advanced Ovarian Cancer. N Engl J Med. 2018;379:2495–505.

26. Tutt ANJ, Garber JE, Kaufman B, Viale G, Fumagalli D, Rastogi P, et al. Adjuvant Olaparib for Patients with BRCA1- or BRCA2-Mutated Breast Cancer. N Engl J Med. 2021;384:2394–405.

27. Li K, Luo H, Huang L, Luo H, Zhu X. Microsatellite instability: a review of what the oncologist should know. Cancer Cell Int. 2020;20:16.

28. Dedeurwaerdere F, Claes KB, Van Dorpe J, Rottiers I, Van der Meulen J, Breyne J, et al. Comparison of microsatellite instability detection by immunohistochemistry and molecular techniques in colorectal and endometrial cancer. Sci Rep. 2021;11:12880.

29. Stewart MD, Merino Vega D, Arend RC, Baden JF, Barbash O, Beaubier N, et al. Homologous Recombination Deficiency: Concepts, Definitions, and Assays. Oncologist. 2022;27:167–74.

30. Fu Y, Qi L, Guo W, Jin L, Song K, You T, et al. A qualitative transcriptional signature for predicting microsatellite instability status of right-sided colon cancer [Internet]. Research Square. Research Square; 2019. Available from: 10.21203/rs.2.10681/v1

31. Ziegler J, Hechtman JF, Rana S, Ptashkin RN, Jayakumaran G, Middha S, et al. A deep multiple instance learning framework improves microsatellite instability detection from tumor next generation sequencing. Nat Commun. 2025;16:136.

32. Anaya J, Sidhom J-W, Mahmood F, Baras AS. Multiple-instance learning of somatic mutations for the classification of tumour type and the prediction of microsatellite status. Nat Biomed Eng. 2024;8:57–67.

33. Davies H, Glodzik D, Morganella S, Yates LR, Staaf J, Zou X, et al. HRDetect is a predictor of BRCA1 and BRCA2 deficiency based on mutational signatures. Nat Med. 2017;23:517–25.

34. Nguyen L, W M Martens J, Van Hoeck A, Cuppen E. Pan-cancer landscape of homologous recombination deficiency. Nat Commun. 2020;11:5584.

35. Abbasi A, Steele CD, Bergstrom EN, Khandekar A, Farswan A, McKay RR, et al. Data from HRProfiler detects homologous recombination deficiency in breast and ovarian cancers using whole-genome and whole-exome sequencing data [Internet]. American Association for Cancer Research. 2025. Available from: 10.1158/0008-5472.c.7906454

36. Gulhan DC, Lee JJ-K, Melloni GEM, Cortés-Ciriano I, Park PJ. Detecting the mutational signature of homologous recombination deficiency in clinical samples. Nat Genet. 2019;51:912–9.

37. Singh AK, Olsen MF, Lavik LAS, Vold T, Drabløs F, Sjursen W. Detecting copy number variation in next generation sequencing data from diagnostic gene panels. BMC Med Genomics. 2021;14:214.

38. Chandramohan R, Reuther J, Gandhi I, Voicu H, Alvarez KR, Plon SE, et al. A validation framework for somatic copy number detection in targeted sequencing panels. J Mol Diagn. 2022;24:760–74.

39. Sanjaya P, Maljanen K, Katainen R, Waszak SM, Genomics England Research Consortium, Aaltonen LA, et al. Mutation-Attention (MuAt): deep representation learning of somatic mutations for tumour typing and subtyping. Genome Med. 2023;15:47.

40. The A, Project G. AACR Project GENIE: Powering Precision Medicine Through An International Consortium, Cancer Discov. 2017;7:818–31.

41. Wen H, Feng Z, Ma Y, Liu R, Ou Q, Guo Q, et al. Homologous recombination deficiency in diverse cancer types and its correlation with platinum chemotherapy efficiency in ovarian cancer. BMC Cancer. 2022;22:550.

42. Cancer Genome Atlas Network. Comprehensive molecular characterization of human colon and rectal cancer. Nature. 2012;487:330–7.

43. Kautto EA, Bonneville R, Miya J, Yu L, Krook MA, Reeser JW, et al. Performance evaluation for rapid detection of pan-cancer microsatellite instability with MANTIS. Oncotarget. 2017;8:7452–63.

44. Niu B, Ye K, Zhang Q, Lu C, Xie M, McLellan MD, et al. MSIsensor: microsatellite instability detection using paired tumor-normal sequence data. Bioinformatics. 2014;30:1015–6.

45. Sztupinszki Z, Diossy M, Krzystanek M, Reiniger L, Csabai I, Favero F, et al. Migrating the SNP array-based homologous recombination deficiency measures to next generation sequencing data of breast cancer. NPJ Breast Cancer. 2018;4:16.

46. Abkevich V, Timms KM, Hennessy BT, Potter J, Carey MS, Meyer LA, et al. Patterns of genomic loss of heterozygosity predict homologous recombination repair defects in epithelial ovarian cancer. Br J Cancer. 2012;107:1776–82.

47. Birkbak NJ, Wang ZC, Kim J-Y, Eklund AC, Li Q, Tian R, et al. Telomeric allelic imbalance indicates defective DNA repair and sensitivity to DNA-damaging agents. Cancer Discov. 2012;2:366–75.

48. Popova T, Manié E, Rieunier G, Caux-Moncoutier V, Tirapo C, Dubois T, et al. Ploidy and large-scale genomic instability consistently identify basal-like breast carcinomas with BRCA1/2 inactivation. Cancer Res. 2012;72:5454–62.

49. Rempel E, Kluck K, Beck S, Ourailidis I, Kazdal D, Neumann O, et al. Pan-cancer analysis of genomic scar patterns caused by homologous repair deficiency (HRD). NPJ Precis Oncol. 2022;6:36.

50. Cosmic. COSMIC [Internet]. 2020 [cited 2024 Jul 30]. Available from: https://cancer.sanger.ac.uk/signatures/id/

51. Howard FM, Dolezal J, Kochanny S, Schulte J, Chen H, Heij L, et al. The impact of site-specific digital histology signatures on deep learning model accuracy and bias. Nat Commun. 2021;12:4423.

52. Ilse M, Tomczak J, Welling M. Attention-based Deep Multiple Instance Learning. In: Dy J, Krause A, editors. Proceedings of the 35th International Conference on Machine Learning. PMLR; 10--15 Jul 2018. p. 2127–36.

53. Devlin J, Chang M-W, Lee K, Toutanova K. BERT: Pre-training of deep bidirectional Transformers for language understanding [Internet]. arXiv [cs.CL]. 2018. Available from: http://arxiv.org/abs/1810.04805

54. Poli M, Massaroli S, Nguyen E, Fu DY, Dao T, Baccus S, et al. Hyena hierarchy: Towards larger convolutional language models [Internet]. arXiv [cs.LG]. 2023. Available from: http://arxiv.org/abs/2302.10866

55. Islam SMA, Díaz-Gay M, Wu Y, Barnes M, Vangara R, Bergstrom EN, et al. Uncovering novel mutational signatures by de novo extraction with SigProfilerExtractor. Cell Genom. 2022;2:None.

56. Sundararajan M, Taly A, Yan Q. Axiomatic attribution for deep networks [Internet]. arXiv [cs.LG]. 2017. Available from: http://arxiv.org/abs/1703.01365

57. Jönsson G, Naylor TL, Vallon-Christersson J, Staaf J, Huang J, Ward MR, et al. Distinct genomic profiles in hereditary breast tumors identified by array-based comparative genomic hybridization. Cancer Res. 2005;65:7612–21.

58. Probabilistic Outputs for Support Vector Machines and Comparisons to Regularized Likelihood Methods.

59. Cortes-Ciriano I, Lee S, Park W-Y, Kim T-M, Park PJ. A molecular portrait of microsatellite instability across multiple cancers. Nat Commun. 2017;8:15180.

60. Dámaso E, Castillejo A, Arias MDM, Canet-Hermida J, Navarro M, Del Valle J, et al. Primary constitutional MLH1 epimutations: a focal epigenetic event. Br J Cancer. 2018;119:978–87.

61. Aaltonen LA, Peltomäki P, Leach FS, Sistonen P, Pylkkänen L, Mecklin JP, et al. Clues to the pathogenesis of familial colorectal cancer. Science. 1993;260:812–6.

62. Guo Q, Househam J, Lakatos E, Nowinski S, Al Bakir I, Grant H, et al. Long deletion signatures in repetitive genomic regions track somatic evolution and enable sensitive detection of microsatellite instability [Internet]. Bioinformatics. bioRxiv; 2024. Available from: https://www.biorxiv.org/content/10.1101/2024.10.03.616572v1

63. Meier B, Volkova NV, Hong Y, Schofield P, Campbell PJ, Gerstung M, et al. Mutational signatures of DNA mismatch repair deficiency in C. elegans and human cancers. Genome Res. 2018;28:666–75.

64. TUMOR TYPES BIOMARKERS FDA-APPROVED THERAPY. Table 1: Companion diagnostic indications [Internet]. [cited 2024 Nov 15]. Available from: https://www.foundationmedicine.com/sites/default/files/media/documents/2024-07/F1CDxTechnical_Specifications_Commercial_SPEC-01197_V3.0.pdf

65. Takeda M, Takahama T, Sakai K, Shimizu S, Watanabe S, Kawakami H, et al. Clinical application of the FoundationOne CDx assay to therapeutic decision-making for patients with advanced solid tumors. Oncologist. 2021;26:e588–96.

66. TruSight Oncology 500 [Internet]. [cited 2024 Nov 15]. Available from: https://emea.illumina.com/products/by-type/clinical-research-products/trusight-oncology-500.html#tabs-48377e6865-item-d4e16e33c1-documentation

67. Zhao C, Jiang T, Hyun Ju J, Zhang S, Tao J, Fu Y, et al. TruSight Oncology 500: Enabling Comprehensive Genomic Profiling and Biomarker Reporting with Targeted Sequencing [Internet]. Bioinformatics. bioRxiv; 2020. Available from: https://www.biorxiv.org/content/10.1101/2020.10.21.349100v1

68. Sinicrope FA, Sargent DJ. Molecular pathways: microsatellite instability in colorectal cancer: prognostic, predictive, and therapeutic implications. Clin Cancer Res. 2012;18:1506–12.

69. Black SJ, Ozdemir AY, Kashkina E, Kent T, Rusanov T, Ristic D, et al. Publisher Correction: Molecular basis of microhomology-mediated end-joining by purified full-length Polθ. Nat Commun. 2020;11:1831.

70. Bennardo N, Cheng A, Huang N, Stark JM. Alternative-NHEJ is a mechanistically distinct pathway of mammalian chromosome break repair. PLoS Genet. 2008;4:e1000110.

71. Beroukhim R, Mermel CH, Porter D, Wei G, Raychaudhuri S, Donovan J, et al. The landscape of somatic copy-number alteration across human cancers. Nature. 2010;463:899–905.

72. Witz A, Dardare J, Betz M, Michel C, Husson M, Gilson P, et al. Homologous recombination deficiency (HRD) testing landscape: clinical applications and technical validation for routine diagnostics. Biomark Res. 2025;13:31.

73. Bergstrom EN, Huang MN, Mahto U, Barnes M, Stratton MR, Rozen SG, et al. SigProfilerMatrixGenerator: a tool for visualizing and exploring patterns of small mutational events. BMC Genomics. 2019;20:685.

74. Díaz-Gay M, Vangara R, Barnes M, Wang X, Islam SMA, Vermes I, et al. Assigning mutational signatures to individual samples and individual somatic mutations with SigProfilerAssignment. Bioinformatics. 2023;39:btad756.

75. Gélard M, Benkirane H, Pierrot T, Richard G, Cournède P-H. Bimodal masked language modeling for bulk RNA-seq and DNA methylation representation learning [Internet]. bioRxiv. 2025. Available from: 10.1101/2025.06.25.661237

